# ERK3/MAPK6 promotes triple-negative breast cancer progression through collective migration and EMT plasticity

**DOI:** 10.1101/2024.06.20.599916

**Authors:** Sofia Morazzo, Soraia Fernandes, Marina Fortea, Helena Skálová, Marco Cassani, Kamila Vrzalová, Filip Kafka, Jan Vrbský, Daniel Pereira de Sousa, Veronika Bosáková, Jaeyoung Shin, Jan Fric, Kristina Haase, Giancarlo Forte

## Abstract

Triple-negative breast cancer (TNBC) is the most aggressive subtype of breast cancer and is associated with high cell plasticity, recurrence, and metastatic rate. During epithelial-to-mesenchymal transition (EMT), cancer cells display EMT plasticity, or partial-EMT features, which are required for breast cancer metastasis, such as collective migration. ERK3 has been implicated in promoting migration and invasion of breast cancer, but the mechanisms remain elusive. Here, we investigated ERK3 expression across patient-derived datasets of breast cancer and established its association with aggressive breast cancer phenotypes and poor clinical outcomes. Leveraging the hypothesis that ERK3 contributes to TNBC progression by supporting a partial-EMT state, we showed that ERK3 is essential in different steps of the metastatic process, especially by enabling collective migration but also by modulating cell-extracellular matrix adhesion, anchorage-independent growth, extravasation and colonization. In conclusion, our results demonstrate that ERK3 contributes to TNBC progression and potentially metastasis by promoting EMT plasticity and collective migration.

## 1. Introduction

Triple-negative breast cancer (TNBC) is the most aggressive subtype of breast cancer (BC). Chemotherapy remains the first-line — and largely the only — treatment available, but overall TNBC presents poor prognosis due to high recurrence and metastasis[1]. These aspects are strongly correlated with the high cell plasticity of TNBC, which is associated with the appearance of cancer stem cells (CSC) and cellular epithelial-to-mesenchymal transition (EMT) features. EMT refers to the process of cellular transition from epithelial (E) to mesenchymal (M) state[2,3]. However, EMT can also encompass intermediate states, referred as hybrid E/M or partial-EMT, where cells display high EMT plasticity[2]. A consensus from the EMT International Association suggests that hybrid E/M states should be defined as epithelial-mesenchymal plasticity (EMP), and are characterized by alterations occurring in several molecular markers, such as E-cadherin, cytokeratins (KRT8, KRT18, KRT5, KRT14), SNAIL, and the integrin ITGB3 (CD61), in combination with morphological and/or behavioural changes. These include loss of apical-basal polarity, decreased cell-extracellular matrix (ECM) adhesion, or gain of the ability to migrate and/or invade surrounding tissue[3,4]. In particular, EMP is required for BC cells to metastasize by promoting collective migration through a basal program[2,5]. This basal program is characterized by KRT14 enrichment in leader cells, and is strongly associated with high metastatic burden in *in vivo* models[5].

The mitogen-activated protein kinases (MAPK), such as the classical MAPKs – ERK1/2, ERK5, p38 isoforms and the c-Jun N-Terminal Kinase (JNK) isoforms -, have been strongly implicated in cancer progression and EMT, including BC[6–9]. The atypical ERK3, coded by MAPK6 gene, is constitutively active, in physiological conditions with a high turnover regulated by degradation, indicating that many functions are based on protein abundance[10]. ERK3 has a pro-tumorigenic role in several types of cancer, such as non-small cell lung cancer[11], prostate, ovarian, cervical and gastric cancers[11–15]. In BC, ERK3 is essential for filopodia formation and modulates changes in cell body area leading to more efficient cancer cell migration and invasion capacity[13,16]. Additionally, ERK3 was shown to be involved in the formation of lung metastasis in *in vivo* models of TNBC[12,14]. However, the mechanism through which ERK3 leads to high metastatic burden is still undisclosed.

In this study, we investigated the role of ERK3 in TNBC progression, focusing particularly on cancer cell migration and metastasis. First, we assessed ERK3 expression changes across BC from patient-derived datasets, showing that its overexpression correlates with clinical parameters, such as tumor subtype, tumor grading and survival metrics. We then demonstrated, using four stable cell lines in which we silenced ERK3, that ERK3 contributes to EMP, by i) enhancing the expression of key molecular markers such as SNAIL and KRT14, ii) promoting pro-metastatic pathways such as Wnt/β-catenin, and YAP/CYR61 and iii) enhancing collective migration of TNBC cells. Finally, we confirmed the role of ERK3 in the metastatic capacity of TNBC cells using advanced 3D models, including a 3D-microphysiological system (3D-MPS) to mimic lung capillaries and a co-culture system of TNBC spheroids with human induced pluripotent stem cell (hiPSC)-derived lung organoids.

## 2. Materials and Methods

### 2.1. Patient-derived Database Analysis

Patient-derived data regarding ERK3/MAPK6 gene expression in normal breast tissue, breast invasive carcinoma or metastatic breast invasive carcinoma was retrieved through user-friendly platforms. Specifically, datasets from the Gene Expression Omnibus of National Center for Biotechnology Information (NCBI-GEO)[17], Genotype-Tissue Expression (GETx)[18] and the Cancer Genome Atlas Pan-cancer (TCGA)[19] and the Therapeutically Applicable Research to Generate Effective Treatments (TARGET)[20] were accessed through the Tumor, Normal and Metastatic tissue plotter (TNMplot; https://tnmplot.com/)[21]. Also, ERK3/MAPK6 gene expression across the different subtypes of breast cancer and between different tumor grades from the NCBI-GEO[17] datasets, Affymetrix U133A or U133Plus2, were accessed through the Gene Expression database in Normal and Tumor tissues platform (GENT2; http://gent2.appex.kr/gent2/)[22]. Patient survival curves to either high or low ERK3/MAPK6 expression, including overall patient survival (OS), recurrence-free survival (RFS) and distant-metastasis free survival (DMFS) were accessed and analyzed in the Kaplan-Meier plotter (https://kmplot.com/analysis/)[23], which derives survival data from several datasets deposited in the European Genome-phenome Archive (EGA, https://ega-archive.org/), NCBI-GEO[17] and from the TCGA project[19].

### 2.2. Cell Culture

Human immortalized breast cancer cell line MDA-MB231 was purchased from the American Type Culture Collection (ATCC® CRM-HTB26™) and cultured in Dulbecco’s Modified Eagle Medium (DMEM) high glucose media from Biosera, supplemented with 10% fetal bovine serum (FBS), 1% penicillin-streptomycin and 1% L-glutamine, purchased from Serana. Human umbilical vein endothelial cells (HUVEC; C2519A) and normal human lung fibroblasts (nhLF; CC-2512) were purchased from Lonza and cultured in coated flasks with 50 μg/ml rat tail collagen I (Roche), with Vasculife (LL-0003) or Fibrolife (LL-0011), respectively, from Lifeline cell systems. All cell cultures were confirmed mycoplasma-free with MycoAlert™ Mycoplasma Detection Kit (Lonza), using luminometer Centro LB960, from Berthold Technologies.

Human iPSC (WiCell, DF19–9–7T)-derived lung organoids (LO) were cultured, differentiated and characterized as previously described [24–27]. They were embedded in Cultrex reduced growth factor, basement membrane, type 2 (∼4mg/ml, Cultrex, RGF BME, Type2; R&D Systems) and maintained in LO complete media which consists of LO basic media -Advanced DMEM/F12 media (Thermo Fisher), with HEPES (15mM, Sigma), penicillin/streptavidin (500 U/ml), GlutaMAX (2.5 mM, Thermo Fisher), N2 and B27 supplement (Thermo Fisher) – which was supplemented with HyClone^TM^ FBS (1%, Fisher Scientific) and FGF-10 (500ng/ml, R&D Systems) before each use. Media was changed twice a week and LOs were used between 70 and 120 days in culture.

### 2.3. Induction of transient and stable knockdown

Transient knockdown of ERK3 was induced by small interfering RNA (siRNA) transfection using 20nM of targeted siRNA (Silencer^TM^ Select, assay ID: 142309) as well as 20nM of non-targeting control (Silencer^TM^ Select Negative Control), both purchased from Thermo Fisher, with SAINT-sRNA transfection reagent (SR-2003-01, Synvolux) according to manufacturer’s protocol, in antibiotic-free conditions for 24 hours. Assays were performed 48 hours post-transfection. The efficiency of each transfection was evaluated by western blotting, as described in section 2.7.

The generation of stable ERK3 knockdown cell lines was obtained using lentiviral packaged short-hairpin RNA (shRNA) particles with green fluorescent protein (GFP) tag and puromycin resistance (Vector Builder). Lentiviral titration of three different targeting shRNAs were tested paired with a scramble control, and shRNA targeting sequence ATCCTTACATGAGCATATATT (MOI: 8 viral particles/cell) was selected to generate the stable cell lines based on knockdown efficiency and viability. Cells were transduced in antibiotic-free media with 8µg/ml of polybrene (Santa Cruz), and selected with puromycin (1µg/ml, Invivogen). A total of 4 different transduced cell lines were generated with ERK3 knockdown (shERK3) and one with the scramble control (shWT). The efficiency of knockdown was evaluated by western blot and real-time PCR (qPCR), as described in section 2.7 and 2.8, respectively.

### 2.4. Migration and Invasion assays

Transwell migration assay was performed using the colorimetric QCM Chemotaxis Cell Migration Assay (ECM508, Merck), in 24 well-plate format with 8µm pore size. Following the manufacturer’s protocol, at 48hours post siRNA transfection 2,5×10^5^ cells/ml were plated in serum-free conditions (300µl) in the top insert and with full media (300µl supplemented with 10%FBS) in the bottom well. After 24 hours, non-migrating cells were scraped out by cell scraper (Corning) and, using the kit’s extraction buffer and cell stain, the remaining attached and migrating cells were solubilized, extracted stained and measured, in a 96 well-plate, by spectrophotometry, at 560nm, using microplate reader Multiskan Go (ThermoFisher).

Wound healing assays were performed with either siRNA transfected cells or shRNA transduced stable cell lines, in 6 well-plate format. Wound area was created by scratching with a 200µl tip, when cell culture reached approximately 70-80% confluency. Two images, of the wound area were acquired per well (2-3 wells/condition) and per time point (0h, 20h), using contrast microscope Leica DM IL Inverted Phase Contrast Microscope and analyzed using ImageJ software and wound healing size tool plugin[28].

3D breast cancer spheroids (BCS) were generated using 2×10^5^ cells/ml, with 0,05mg/ml collagen I solution, rat tail (Gibco A10483), plated in ultra-low attachment, round bottom 96 well-plates and centrifuged at 1000RPM for 5 minutes, with 5 acceleration and 2 deceleration speed. After 24h-48h, spheroids were completely formed and used either for the invasion assays or confrontation assays. For the invasion assays, the spheroids were embedded in either approximately 4mg/ml Cultrex, (RGF BME, Type2, R&D Systems) or in 1 mg/ml collagen I solution, rat tail (Gibco A10483), in 4 well-plate. Images were acquired using contrast microscope Leica DM IL Inverted Phase Contrast Microscope and analyzed using ImageJ software, with manual selection tool, to measure invading area at different time points (day 0, day6). Their use in the confrontation assays is described below.

### 2.5. Cell-ECM or cell-dECM adhesion assays

Adhesion assay to different recombinant ECM proteins was performed using colorimetric ECM Cell Adhesion Array Kit (ECM540, Merck) plating 3×10^4^ cells/well at 48h post-siRNA-transfection. After 24-hour incubation, following manufacturer’s instructions, wells were washed with PBS three times, and, using the kit’s extraction buffer and cell stain, the remaining attached cells were solubilized, extracted, stained and measured, in a 96 well-plate, by spectrophotometry, at 560nm, using microplate reader Multiskan Go (ThermoFisher).

Decellularized ECM adhesion (dECM) assays were performed using either lung dECM hydrogel kit (CC175, Sigma-Aldrich) or liver dECM hydrogel kit (CC174, Sigma-Aldrich), according to manufacturer’s instructions. Briefly, the hydrogels were prepared as 3mg/ml and used to coat 96 well-plate for fluorometric measurement (black, transparent bottom). Stable cell lines were plated at 3×10^4^ cells/well and after 24-hour incubation, wells were washed three times with PBS and remaining cells were measured, based on a standard curve and GFP fluorescence with Biotek FLx800 Fluorescence Microplate Reader.

### 2.6. Soft-agar colony formation Assay

Anchorage-independent growth was evaluated using CytoSelect™ 96-Well Cell Transformation Assay (CBA-130, CellBioLabs), following manufacturer’s instructions. Control and stable ERK3 KD cell lines were plated at 1,5 × 10^3^ cells/well. After 8 days in culture, cells were solubilized, lysed and stained with CyQuant® GR Dye, according to manufacturer’s protocol and measured by standard curve and fluorometric detection with Biotek FLx800 Fluorescence Microplate Reader.

### 2.7. Western blot

Proteins were extracted from cell pellet with RIPA buffer (NaCl 150mM, 50mM Tris, 1% Nonidet P-40, 0.5% sodium deoxycholate, pH 7.4), supplemented with PMSF 10mM, protease inhibitor and phosphatase inhibitor cocktails (1% v/v, P8340 and P5726, respectively, Sigma-Aldrich) and 1% SDS. Protein amount was determined using Pierce BCA Protein Assay kit (Thermo Fisher), using albumin to generate a 2-fold dilution standard curve and measuring by spectrophotometry (562nm). Samples were then prepared as 8µg total protein, together with 4x Laemmli Sample Buffer (Biorad) and 1mM DTT, and denatured for 10min at 95°C. They were loaded in 10% acrylamide gels (TGX Stain-Free™ FastCast™ Acrylamide Starter Kit, 10%, Biorad), with protein ladder (NZYColour Protein Marker II, NZYtech). Densitometric data was obtained by incubating with Clarity™ Western ECL Substrate (Bio-Rad), for 1 minute, and acquisition in ChemiDoc MP Imaging System (Bio-Rad) using ImageLab 6.0.1 software (Biorad). Using Stain-Free technology, loading and transfer efficiency to nitrocellulose membrane was visually evaluated. Densiometric data was normalized against anti-beta-tubulin (1:5000, HRP-conjugated, ab21058, Abcam), or total protein obtained from Stain-Free images.

Primary antibodies used were anti-ERK3 (1:500, Mouse mAb M02, clone 4C11) from Abnova, anti-GFP (1:1000, Mouse mAb sc-9996) from Santa Cruz Biotechnology and anti-cytokeratin 14 (1:500, ab181595 [EPR17350], rabbit), from Abcam. Also, anti-SNAIL (1:1000, C15D3 Rabbit mAb #3879), anti-SLUG (1:1000, C19G7 Rabbit mAb #9585), anti-N-cadherin (1:1000, D4R1H XP® Rabbit mAb 13116), anti-YAP (1:1000, D8H1X XP ® Rabbit mAb #1474), anti-CYR61 (1:1000, D4H5D XP ® Rabbit mAb, #14479), anti-β-Catenin (1:1000, D10A8 XP® Rabbit mAb #8480); purchased from Cell Signaling. Secondary antibodies used were HRP-conjugated anti-rabbit (1:2000) and anti-mouse (1:2000), from Cell Signaling.

### 2.8. cDNA sample preparation and qPCR

RNA was extracted from cell pellet, using High Pure RNA Isolation Kit (Roche), according to manufacturer’s protocol, and cDNA was synthetized using Transcriptor First Strand cDNA Synthesis Kit (Roche), according to manufacturer’s protocol. qPCR was performed using LightCycler 480 SYBR Green I Master (Roche) and StepOnePlus™ Real-Time PCR System (Thermo Fisher), according to manufacturer’s protocol, with the primers listed in table 1. Gene expression was normalized to beta-actin (ACTB).

**Table 1.**
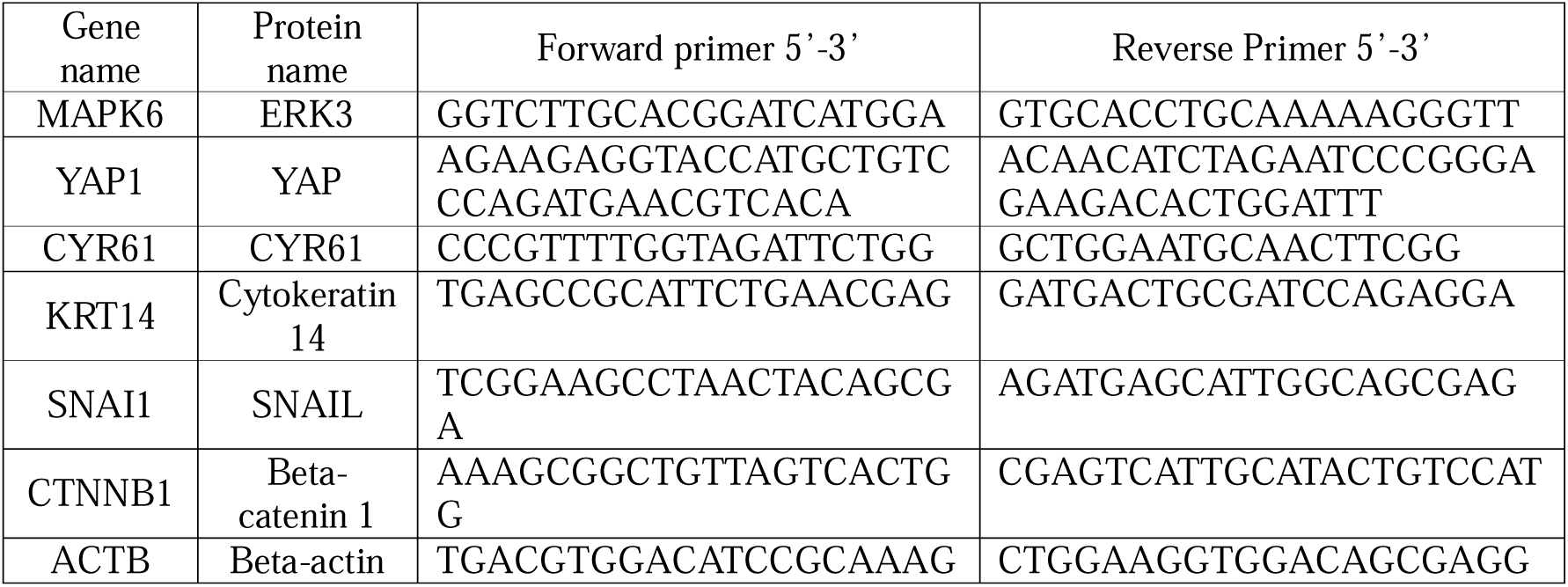
qPCR primers for gene expression determination.

### 2.9. RT-Profiler

Gene expression array was performed using RT² Profiler™ PCR Array Human Wound Healing Profiler PCR mRNA array, for wound healing markers (PAHS-121Z, QIAGEN), following manufacturer’s instructions, in LightCycler Galaxy (Roche). Gene expression data was analyzed in GeneGlobe Analyzer from QIAGEN (https://geneglobe.qiagen.com/cz/analyze)

### 2.10. Device Fabrication

For the extravasation assay, devices were fabricated as previously described[29], mixing in a 10:1 proportion PDMS (SYLGARDTM 184 Silicone Elastomer Kit, Dow) with elastomer, following the manufacturer’s protocol. After mixing, it was degassed, poured into the respective mould, degassed again and cured at 60°C overnight. Then, from the mould, devices were cut out and biopsy punches of 1 and 2mm diameter were used to create the entry to the central channel and the media ports, respectively. The devices were then air-plasma bonded to disinfected glass slides and incubated again at 60°C overnight to restore their native hydrophobic state. Before seeding, devices were sterilized in laminar flow hood under ultraviolet light for at least 30 minutes.

### 2.11. Microvasculature Formation

Following the previously described protocol[29] to generate perfusable vasculature with the above-mentioned device, fibrinogen, derived from bovine plasma (F8630, Sigma), solution should be prepared ahead of time to 12mg/ml diluted in phosphate-buffered saline (PBS). Before seeding, fibrinogen solution should be filtered with a 0.2µm pore size filter and thrombin (T4648, Sigma) stock solution (100 U/ml in 0.1% w/v bovine serum albumin solution) should be diluted to a final concentration of 4U/ml in cold VascuLife media. After trypsinization with TryplE, HUVECs (24×106 cells/ml) and nhLF (4.8×106, cells/ml) are mixed individually in thrombin solution Each cell-thrombin suspension is then mixed together in a 1:1 ratio and with fibrinogen to a final concentration of 3mg/ml. The cell-fibrinogen mixture (15-20µl) is then injected in the central channel of the device and incubated for approximately 20 minutes, in a humidity chamber to allow the gel to polymerize. After, VascuLife media (150µl) is added to the media ports and devices are maintained in a large Petri dish (150mm diameter), to reduce contamination, and with a smaller Petri dish (30mm diameter) filled with PBS, to avoid dehydration. Media was changed daily.

### 2.12. Extravasation Assay

After 7 days in culture, cancer cells were injected into the devices. Right before adding the cancer cells, the devices were first incubated with 0.1% bovine serum albumin (BSA) in PBS, sterilized with a 0.2µm pore size filter, for 15-20 minutes to avoid unspecific adhesions, as previously described[30]. Stable cell lines with and without ERK3 knockdown with GFP label were trypsinized in TryplE and resuspended in VascuLife with 5% FBS, and approximately 4.6×10^4^ cells were injected through the media ports. Each biological replicate of shERK3 was used in independent rounds, consisting of 6-8 devices per time point (6, 24, 48 and 72 hours), including 2 devices as control per time point (no injection of cancer cells but all other steps equal). A total of 4 rounds were completed, one with each biological replicate. After permeability measurement, devices were washed with PBS and fixed with 4% paraformaldehyde (PFA) for at least 5 hours, then washed three times with PBS and stored in 0.01% sodium azide in PBS, sealed with parafilm, at 4°C.

Then, devices were stained with Wheat Germ Agglutinin, Alexa Fluor™ 647 Conjugate (WGA, Thermo Fisher) overnight at 4°C, and washed the next day, three times in PBS, before imaging. Images were acquired using Zeiss LSM 780 confocal microscope. A central region was selected at random, and using 20x objective with 0.6 zoom, the tilescan function (7×5 tiles) was used to acquire an area representing approximately 50% of the whole device; and the z-stack function (3.6µm step size) with a channel for GFP and T-PMT detection, and a channel for WGA 647. Images were then converted in Imaris File Converter 9.9.1 and processed in Imaris Viewer. Specifically, the background was subtracted and images were aligned, and then using “surface” tool based of the WGA channel, vessel network was determined. Then cancer cells were manually counted and classified, in relation to the vessels. Specifically, if the whole cell body was inside the vascular space, they were labeled as intravascular, and if completely outside the vascular space, they were labeled as extravascular. In cases in which the cell body is in both spaces, they were labeled as in mid-extravasation. Furthermore, many intravascular cells were in clusters, when possible to count 2 or 3 cells, these were considered small clusters, while if 4 cells or more they were considered big clusters.

### 2.13. Permeability Measurement

At each time point (24, 48 and 72 hours), permeability was measured by removing the media and adding to one media channel 40µl of 70kDa FITC-conjugated dextran (46945, Sigma) prepared as 0.1mg/ml in VascuLife. After approximately 1 minute, to allow perfusion, 40µl of the same solution was added to the opposite media channel to stop the flow. Time-lapse images were acquired using Stellaris 8 confocal microscope and LAS X Leica software. Specifically, 10x objective was used, and 4 random central regions of the device were selected to acquire z-stack time-lapse image (3 x 5 min interval), with 5µm step size. Permeability – *P*(cm/s) – was calculated as described in depth here[29], in ImageJ, using maximum projection images from the dextran channel at *t*=0 to generate a binary mask and outline the vessel perimeter and extravascular tissue area.

### 2.14. Confrontation Assay

Human iPSC-derived lung organoids (LO) were co-cultured with BCS generated from shRNA stable cell lines. The LO was placed, embedded in Cultrex RGF BME type 2 (∼4mg/ml, R&D Systems), and three BCS of either shWT or shERK3 were placed around the LO, in proximity, in 4 well-plates, for each biological replicate (shERK3 transduction). After approximately 40 minutes of incubation at 37°C, to allow the cultrex to gelify, LO complete media was added. Images were obtained using Zeiss LSM 780 confocal microscope, detecting GFP signal and T-PMT/Bright Field, with 10x objective, tile scan function and Z-stack (40 stacks), for each time point (day1 – day6 or day 8, every 24 hours). After 6 or 8 days in culture, the confrontation assays were then washed in PBS and fixed with 4% PFA for 45 minutes, then washed three times in PBS and incubated in 15% sucrose with 0,03% eosin at 4°C overnight. The next day, samples were embedded in Optimal cutting temperature compound (OCT) for cryosectioning.

### 2.15. Immunofluorescence

Samples from the confrontation assay were stained with Wheat Germ Agglutinin, Alexa Fluor™ 647 Conjugate (WGA, Thermo Fisher) for 10 minutes, following permeabilization in 0,2% triton for 10 minutes, blocking for 30 minutes in 2,5% bovine serum albumin (BSA) and incubation with anti-GFP conjugated with Alexa Fluor™ 488 (1:300, sc-9996, Santa Cruz Biotechnology) for 1 hour, then DAPI for 15 minutes. After washing with PBS, slides were mounted with Mowiol overnight before acquisition. Autofluorescence was quenched by incubating with PrestoBlue™ Cell Viability Reagent (Thermo Fisher) for 1 minute, as reported here[31–33]. Images were acquired using Zeiss LSM 780 confocal microscope and processed in ImageJ software.

### 2.16. Proliferation Assay

Proliferation activity was measured in stable cell lines, using Click-iT™ Plus EdU Alexa Fluor™ 647 Flow Cytometry Assay Kit (Thermo Fisher), according to manufacturer’s protocol, in Flow Cytometer (FACS Canto II, BD Biosciences) and then analyzed using FlowJo software, version 10.

### 2.17. Statistical Analysis

Experiments were performed either using independent transfections of siRNA or the four different stable cell lines generated by shRNA transduction via lentiviral delivery to assure reproducibility. GraphPad Prism 10 was used to perform statistical analysis by either student t-test, one way-ANOVA followed by Tukey’s multiple comparison test or two-way ANOVA followed by Sídák’s multiple comparisons test. Survival curve analysis by log-rank test was performed in the Kaplan-Meier plotter[23].

## 3. Results

### 3.1. ERK3 is mostly expressed in aggressive subtypes of BC and correlates with poor clinical outcome in patients

The significance of ERK3 in BC was initially evaluated by exploring large multi-omics datasets, including the GETx[18], TCGA[19] and TARGET[20], as well as datasets from the NCBI-GEO[17], through user-friendly platforms such as the TNMplot[21], GENT2[22], and the Kaplan-Meier plotter[23]. ERK3 expression across patient samples was correlated with various clinical parameters.

Firstly, we observed that ERK3, coded by MAPK6, is significantly more expressed in healthy breast tissue samples in comparison with other members of the MAPK family such as ERK1/2 (MAPK3/MAPK1), ERK5 (MAPK7), the p38 isoforms (MAPK11, MAPK12, MAPK13, MAPK14) or the JNK isoforms (MAPK8, MAPK9, MAPK10; Fig. 1A). Moreover, we found that ERK3 is overexpressed in tumor and metastatic BC samples when compared to normal breast tissue (Fig.1A and 1B). Since inter-patient variability can affect gene expression, we then verified ERK3 overexpression in primary tumors by analysing paired samples with normal-adjacent tissue from the same patient (S1). Secondly, given the significant impact of the distinct molecular subtypes of BC on metastatic rate and prognosis, ERK3 expression was further examined, based on the three established clinical groups, estrogen/progesterone receptor positive (ER/PR+), human epidermal growth factor receptor positive (HER2+) and TNBC subtypes[34]. The analysis revealed that ERK3 is significantly more expressed in the most aggressive subtypes, i.e. HER2 and TNBC (Fig. 1C). Thirdly, a key histopathological feature of TNBC, in comparison to the other subtypes, is the higher tumor grading, which refers to less differentiated cells, higher plasticity and more aggressive metastatic potential[35]. Our analysis shows that ERK3 expression correlates with tumor grading being mostly expressed in grade 3 samples, compared to grade 1 or 2 tumors (Fig. 1D).

**Figure 1.**
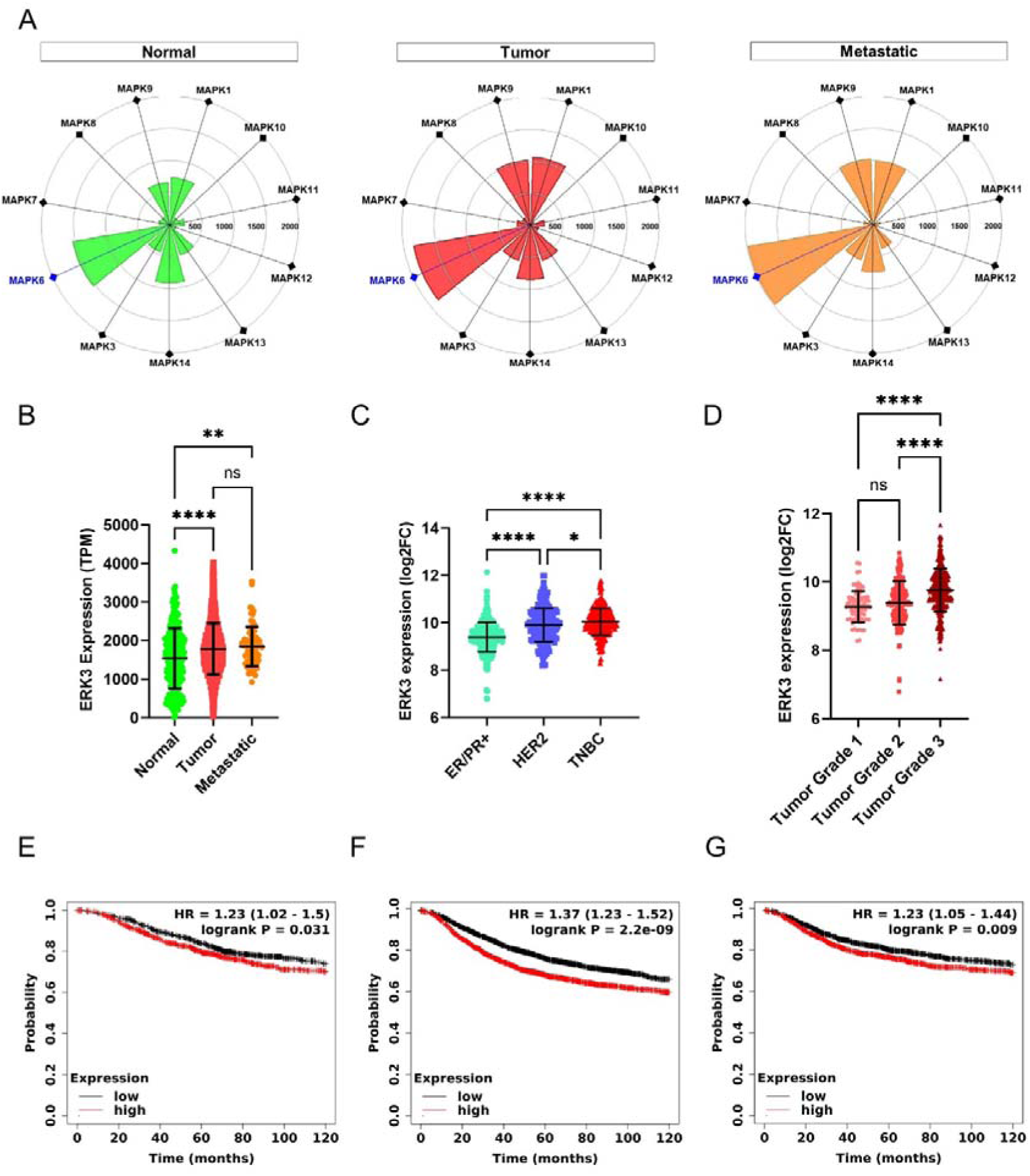
ERK3 overexpression in BC patients correlates with poor patient survival. (**A)** Targetgram analysis of the average expression (transcripts per million, TPM) of the different MAPK genes, in normal breast tissue and in primary and metastatic breast tumors, extracted directly from TNMplot[21]. **(B)** Dotplot representation of ERK3 gene expression (TPM), in normal breast tissue and in primary and metastatic breast tumors (Normal-N=242, Tumor-N=7440, Metastatic-N=77), from the TNMplot. ERK3 gene expression is shown as log2 fold change (log2FC), from the GENT2 platform **(C)** distributed by molecular subtype (ER/PR+ -N=623, HER2 – N=229, TNBC – N=251), and (**D)** by BC histological grade (grade 1 – N=63, grade 2 – N=151, grade 3 – N=358). Survival curves showing the probability of overall survival – (OS)**(E)**, recurrence-free survival (RFS)**(F)** and distant-metastasis free survival (DMFS)**(G)** of BC patients with either high (red) or low (black) expression of ERK3. Accessed and analyzed through the Kaplan-Meier plotter platform. Dotplot graphs show data as mean ± SD. Statistical analysis was performed either by one-way ANOVA followed by Tukey’s multiple comparisons test (B, D and E), paired student t-test (C) or by log-rank test (F, G and H). HR – hazard ratio. * *P* < 0.05, \*\**P*<0.01, \*\*\*\**P*<0.0001 or ns – non-significant.

Finally, TNBC has the worse prognosis with BC patients due to higher recurrence rate, particularly within 5 years of diagnosis[36]. Also, TNBC has a higher metastatic rate and in the surveillance, epidemiology, and end results program (SEER), the 5-year survival rate of localized TNBC drops from an average of 80% to 12% when distant metastasis form[37]. Therefore, we analyzed the impact of ERK3 expression in BC patients not only in overall survival (OS), but also in recurrence-free survival (RFS) and distant-metastasis-free survival (DMFS). Our results indicate that high expression of ERK3 results in decreased OS (Fig. 1E, 115 months for low expression vs 81,6 months for high expression, HR=1.23), RFS (Fig. 1F, 64,9 months for low expression vs 38,4 months for high expression, HR=1.37) and DMFS (Fig. 1G, 100 months for low expression vs 68,4 months for high expression, HR=1.23).

### 3.2. ERK3 promotes collective BC migration and invasion

We next set at investigating whether ERK3 might play a role in collective and single breast cancer migration, in which the former is an indicator of epithelial-mesenchymal plasticity and associated with more aggressive and efficient metastasis of BC and the latter is an indicator of full EMT and associated with cancer recurrence[2]. We transfected MDA-MB231, which is the most aggressive TNBC cell line[38], with siRNA against ERK3 or a scramble control (siERK3 and siWT, respectively; Fig. 2A). Then, we investigated ERK3 effect in single-cell and collective migration, using the transwell and wound healing assays, respectively. For the transwell migration assay, serum-free condition was used as a negative control for chemoattraction compared to full media condition (containing 10%FBS). The obtained results showed no differences in cellular migration rates, indicating that ERK3 does not contribute to single-cell migration in a general chemoattractant setting (Fig. 2B). However, when investigating collective migration using the wound healing assay, we observed that ERK3 knockdown resulted in slower collective migration at 20 hours (shERK3:-16.62% ± 5,451% compared to average siWT; Fig. 2C and 2D). As the migration process is intimately connected with the balance of cell-ECM adhesion,[39] a cell-ECM adhesion assay was performed to further elucidate this result. The data shown in Fig. 2E revealed that ERK3 knockdown significantly increased cell adhesion to various ECM substrates typical of breast tissue, including basement membrane proteins such as collagen type IV and laminin, or interstitial proteins such as collagen type I, fibronectin, tenascin and vitronectin.[40] Together, these results suggest that ERK3 induces a pro-migratory phenotype by reducing the adhesion of breast cancer cells to ECM.

**Figure 2.**
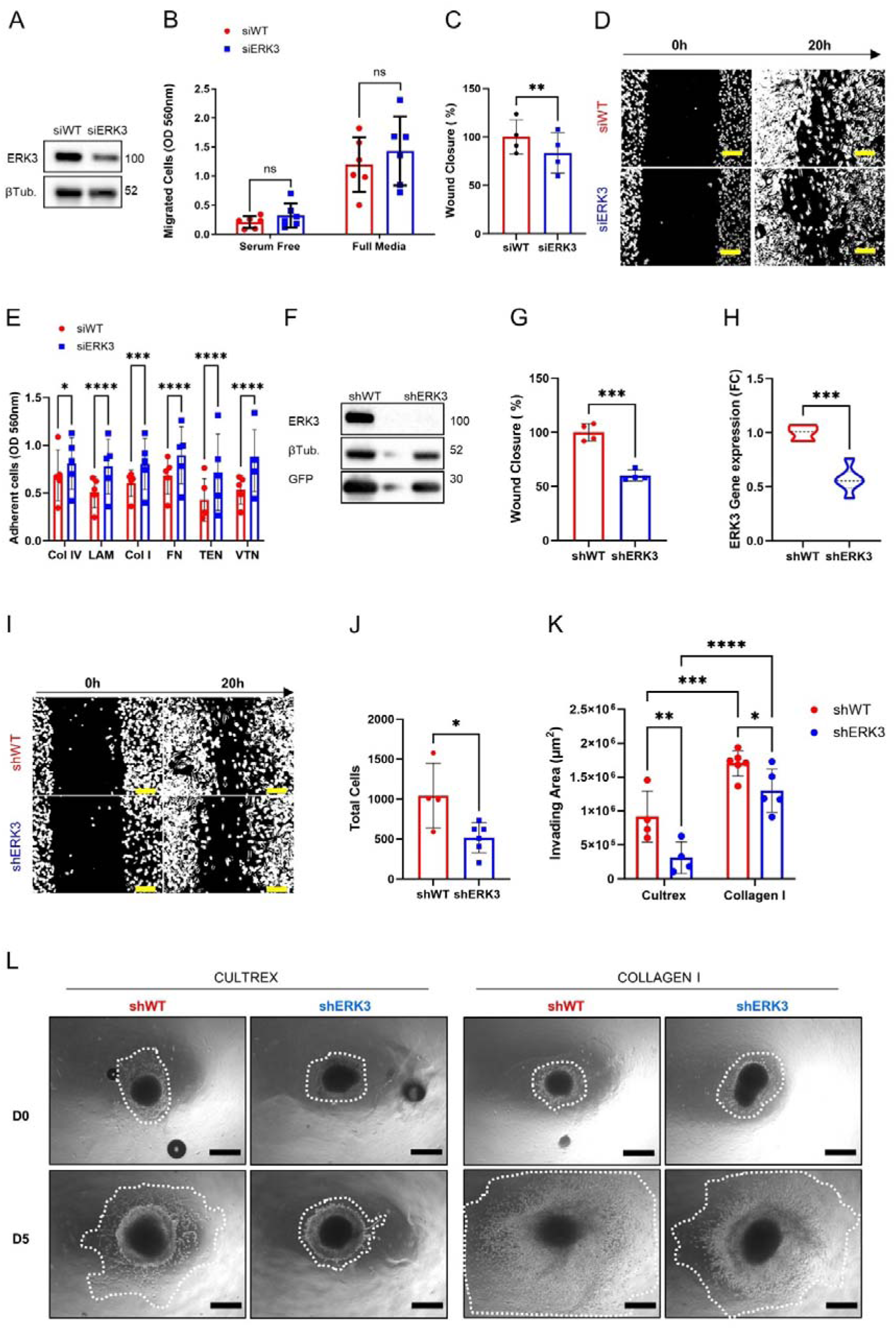
– ERK3 promotes collective migration and invasion of TNBC cells. **(A)** Transient ERK3 knockdown in MDA-MB231 cell line by siRNA and respective scramble control (siERK3 and siWT, respectively) was validated by western blot, for ERK3 and β-tubulin (βTub.). **(B)** Transwell migration assay using siERK3 and siWT cells in either serum-free or full media conditions (N=6 for 20h). **(C)** wound healing assay using siERK3 and siWT cells (N=4) and **(D)** respective representative image of the wound healing assay at 0 h and 20 h (Scale bar: 100µm). **(E)** siERK3 and siWT cells-ECM adhesion assay, across different ECM substrates (Col IV – collagen type IV, LAM – laminin, Col I – collagen type I, FN – fibronectin, TEN – tenascin, VTN – vitronectin; N=5). **(F)** and **(G)** Stable MDA-MB231 cell lines with and without shRNA-induced ERK3 silencing (shERK3 and shWT, respectively), were validated by western blot for ERK3, β-tubulin (βTub.) and GFP **(F)**, and by qPCR for ERK3 gene expression (N=3) **(G)**. (**H)** and **(I)** Wound healing assay performed using shERK3 and shWT stable cell lines (N=4), as characterized by qPCR analysis **(H)**, and representative image of the wound healing assay at 0 h and 20 h **(I)** (Scale bar: 100µm). **(J)** Total cell number determined in soft-agar colony formation assay, after 8 days in culture (shWT -N=4, shERK3 – N=6). **(K)** and **(L)** Migration and invasion ability determined by quantifying the invading area of TNBC spheroids at day 0 and day 5 of culture, in either cultrex or collagen (cultrex -N=4; collagen I – N=6) (K), and representative images with the corresponding invading area **(L)** (white dotted lines; Scale bar: 250µm). All data are presented as mean ± SD. Statistical analysis was performed either by paired student t-test (B and C), unpaired student t-test (F, G and I), or two-way ANOVA followed by by Šídák’s multiple comparisons test (D and J) * *P* < 0.05, \*\**P* <0.01,*** *P* < 0.001, \*\*\*\**P*<0.0001 or ns – non-significant.

To better investigate the role of ERK3 in different metastasis-related processes, we established stable GFP-expressing MDA-MB231 cell lines using 2 independent shRNAs directed against ERK3 (shERK3). Four biological replicates were generated, along with the scramble control (shWT). The stable cell lines were validated by western blot and qPCR to confirm the downregulation of ERK3 (Fig. 2F, 2G and S2A).

We first corroborated the results obtained for the wound healing assay, by assessing cell migration at 0 and 20 hours using the shERK3 and shWT stable lines. The results confirmed that ERK3 affects collective migration as shown by the lower wound closure detected for shERK3 cells in comparison to shWT (shERK3: −39.95%± 4.698% compared to average shWT; Fig. 2H and 2I).

Another process associated with reduced cell-ECM adhesion and enhanced metastasis is anchorage-independent survival[41]. We investigated this phenomenon on our stable cell lines, using the soft-agar colony formation assay, and showed that ERK3 promotes cancer cell ability to survive ECM detachment (Fig. 2J). Next, to confirm that the results obtained from the wound healing and soft-agar colony formation assays were not due to differences in proliferation activity between shERK3 and shWT cells, we used the Edu incorporation assay and demonstrated that ERK3 does not impact the proliferation of these cells (Fig. S2B). This outcome strongly supports the hypothesis that ERK3 modulates collective migration and survival mechanisms independently of breast cancer cell proliferation.

To further reinforce these findings, we used a 3D model in which TNBC spheroids were embedded in either cultrex or collagen type I hydrogels to study the migration and invasion of the cells in the presence or absence of ERK3. Cultrex 3D scaffold mimics the basement membrane of normal tissue, which is usually degraded by invasive BC. During the process, the invasive cells come into contact with collagen type I, an interstitial ECM protein that is abundant in the stroma of breast tissue and is increased in invasive BC tissue[16,40]. The results of our experiment showed that spheroids from cells without ERK3 silencing (shWT) were able to migrate and invade through both substrates more efficiently than those from cells with ERK3 silencing (shERK3), as it can be visible from the measured invading areas (Figure 2K). Moreover, we also observed that the overall migration and invasion ability of shWT and shER3 cells was affected by the ECM substrate used for the experiment, with enhanced migration through collagen I compared to cultrex (Fig. 2K and 2L).

Altogether, these results show that ERK3 promotes EMP features associated with metastasis by supporting collective migration and enhancing survival and invasion capabilities of TNBC cells.

### 3.3. ERK3 upregulates the expression of key EMT and pro-metastatic markers in TNBC cells

Reduced cell—ECM adhesion and collective migration are two functional features characteristic of cells undergoing partial-EMT or EMP[3]. Since ERK3 decreases cell—ECM adhesion and promotes collective migration in TNBC cells (Fig. 2), we set at investigating whether ERK3 facilitates EMP in these cells. Key drivers of EMP in TNBC are the transcription factor SNAIL and β-catenin[42]. Importantly, BC, including TNBC, requires a basal program to promote collective migration and successful metastization, in which there is an enrichment of cytokeratin 14 (KRT14), particularly in leader cells[5,43]. Therefore, we evaluated the expression of these markers, together with other key proteins involved in EMT such as YAP and its downstream effector CYR61, in shERK3 and shWT cells. Western blot analysis, performed with our biological replicates, showed that ERK3 silencing lead to the downregulation of EMP markers SNAIL, β-catenin and KRT14 as well as YAP and CYR61 (Fig. 3A). Furthermore, to investigate if the observed downregulation also occurred at the transcriptional level, we performed qPCR, revealing that ERK3 silencing also reduces the transcription of the encoding genes for SNAIL (SNAI1), β-catenin (CTNNB1), YAP (YAP1) and CYR61 (Fig. 3B).

**Figure 3.**
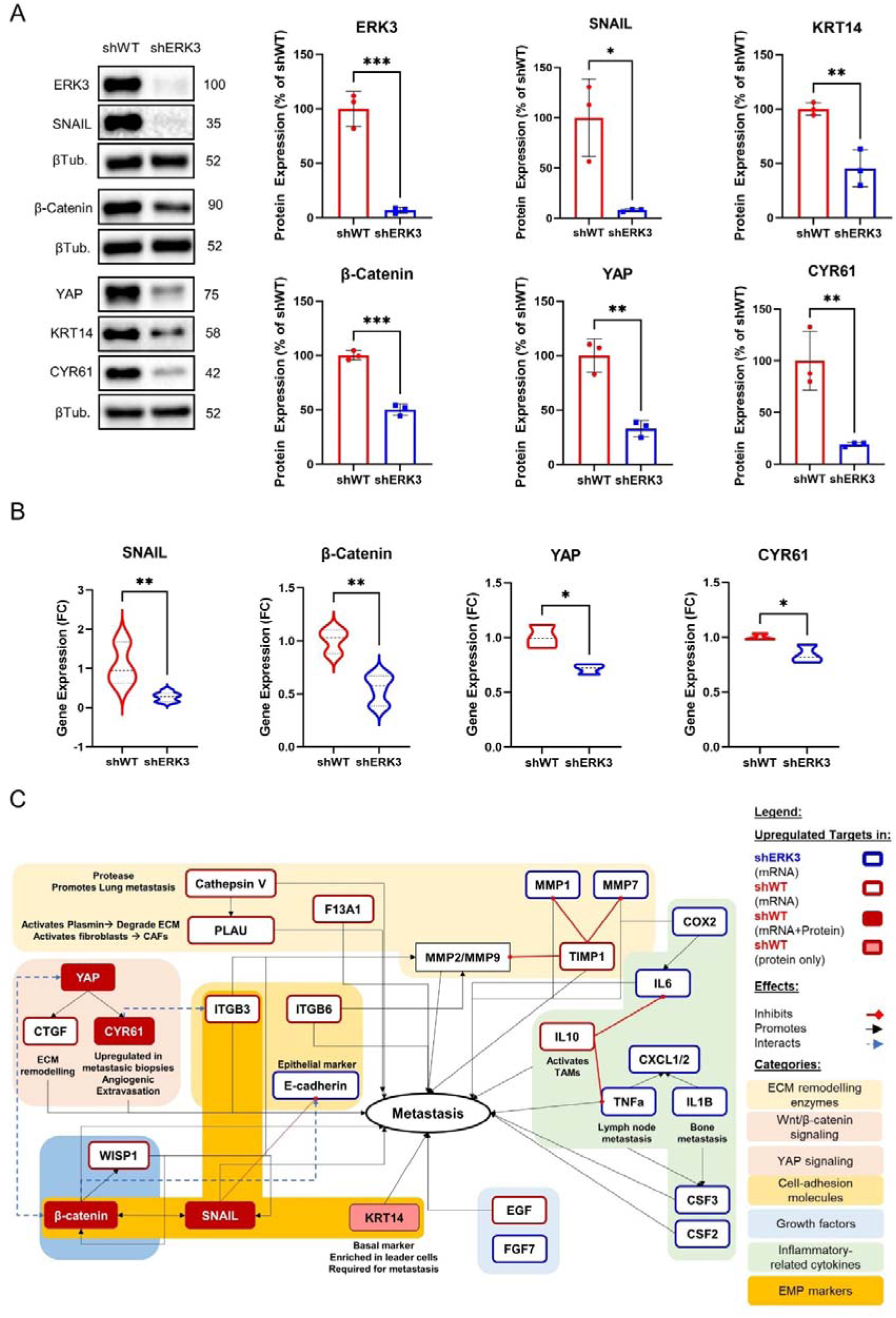
– ERK3 upregulates EMT, pro-metastatic and collective migration markers in TNBC cells. **(A)** Representative images of western blot and relative protein quantification of ERK3, SNAIL, β-catenin, YAP, CYR61, cytokeratin 14 (KRT14), and β-tubulin (βTub.) in shERK3 and shWT stable cell lines. **(B)** Gene expression determined by qPCR and expressed as fold change (FC) of SNAIL, β-catenin, YAP and CYR61 in shERK3 and shWT stable cell lines **(C)** Schematic representation of the differentially regulated markers by ERK3 as determined by RT^2^ profiler PCR Array, western blot and qPCR, and how they relate to the EMT and/or metastatic process in BC, as reported in literature. Data presented mean ± SD. Statistical analysis was performed using unpaired *t-test.* (N=3). *\* P* <0.05*, **P* <0.01, \*\*\**P* <0.001

To gain a better understanding of ERK3 impact on the transcription of other important migration markers, we screened the expression by RT-PCR Array of 84 genes involved in cell migration and invasion (Fig.3C and Table S1). Out of the 84 targets, 25 were differentially regulated in shERK3 and shWT cells. In shERK3 cells, we found the upregulation of 14 genes, which include several inflammatory cytokines (CSF3, IL1B, PTGS2, CXCL11, TNF, CXCL1, IL6, CSF2, and CXCL2), genes related to ECM degradation (MMP1, MMP7, and PLAT), the growth factor FGF7, and notably, the gene encoding E-cadherin (CDH1), a key epithelial marker. In contrast, in the control cells (shWT) there was the upregulation of 11 genes, 10 of which known to promote metastasis in BC. Among these genes, we found coagulation factor XIII (F13A1), which converts fibrinogen to fibrin, leading to the formation of clots and tumor emboli; the powerful anti-inflammatory cytokine IL10; CCN family genes (CCN2/CTGF and CCN4/WISP1); integrins (ITGB3, ITGB6); ECM remodelling genes (CTSV, PLAU, TIMP1); the growth factor and EMT inducer EGF; and the tumor suppressor PTEN (Fig. 3C and Table S1)[41,44–58].

Overall, these findings show that ERK3 expression is linked with the expression of key partial-EMT markers – SNAIL and KRT14 -, and promotes the expression at both transcriptional and protein levels of pro-metastatic markers, i.e. β-catenin, YAP and CYR61.

### 3.4. ERK3 increases cancer cell extravasation capacity

Metastatic cancer cells are characterized by their ability to disseminate towards distant organs, where eventually secondary tumors are established[2]. In this regard, TNBC cells have been shown to preferentially metastasize to the lung [1,34,35]. In order to investigate the role of ERK3 in this process, we used a complex 3D microphysiological system (3D-MPS) in which lung fibroblasts and endothelial cells were seeded within a fibrin gel, creating a perfusable vasculature that mimics the healthy human lung capillaries (Fig. 4A). Using this setup, first, we investigated the permeability of the 3D-MPS (24, 48 and 72 hours post-cancer cell injection) by adding a 70 kDa FITC-conjugated dextran solution to one media port, and after 1 minute adding the dextran solution to the opposite media port, in order to stop the flow and acquiring time-lapse images. No differences were detected in the diffusion of FITC-conjugated dextran between 3D-MPS inoculated with shWT or shERK3 cells nor with a negative control (no injected cancer cells),), and they were all within physiological parameters (S3, approx..10^-7^cm/s), as previously defined[59]. These results confirmed that the vasculature in our system was permeable and perfusable. Subsequently, the same 3D-MPS were used to quantify TNBC cell extravasation at the different time points (24, 48 and 72 hours). In detail, the devices were immediately fixed after permeability measurement and the cells stained with WGA, before confocal microscopy images were obtained and further processed using IMARIS to improve the definition and visualization of TNBC cells relative to the vascular network (Fig.4A). TNBC cells were classified as intravascular (IV -whole cell body is inside the vascular space) or extravascular (EV -whole cell body is outside the vascular space). Cells that were transmigrating to the extravascular space but with part of the cell body still visible in the IV space were classified as mid-extravasated (ME). Additionally, IV cells were subclassified in single IV cells (S-IV), small cluster (2-3 cells, SC-IV) or in a big cluster (4 or more, BC-IV) (Fig. 4B). The quantification of the transmigration process showed an increase in the mid-extravasated cells in the control cells (shWT) compared to shERK3 cells at 72 hours (Fig. 4C). This result supports the evidence of ERK3 playing a pro-metastatic role in TNBC.

**Figure 4.**
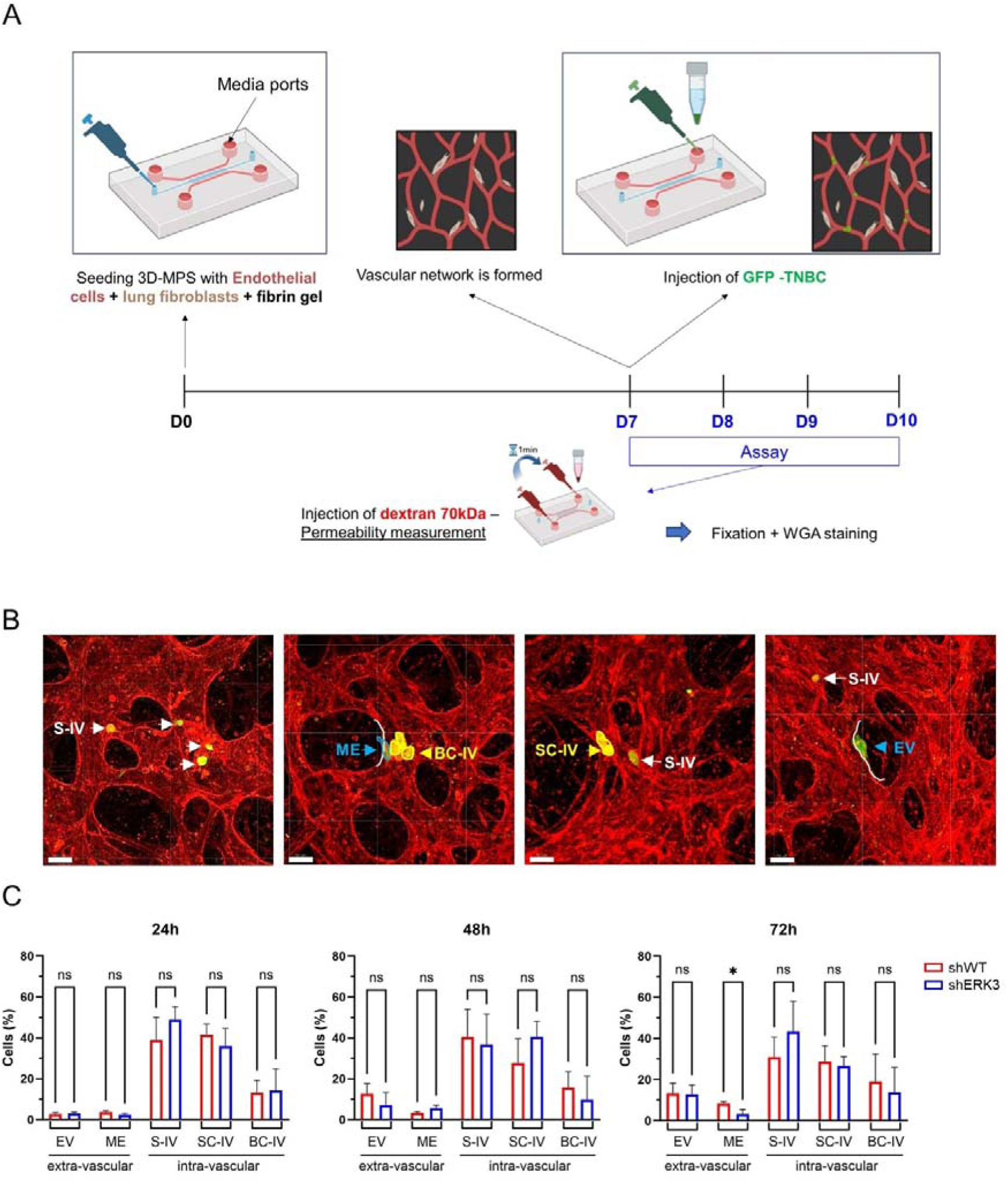
– ERK3 increases cancer cell extravasation in a 3D microphysiological system, which mimics lung capillaries. **(A)** Schematic representation of the 3D-MPS with a central channel (blue) used for the seeding of endothelial cells (red) and lung fibroblasts (brown) within a fibrin gel (top). The parallel media ports used for the injection are indicated by an arrow. Experimental timeline representation of the assay is also shown (bottom). **(B)** Representative confocal images of vasculature stained with WGA (red). GFP-TNBC cells are in green. Depending on their position relative to the blood vessel wall (white line), TNBC cells were classified in intravascular (IV) extravascular (EV) or mid-extravasation (ME, blue). Additionally, depending on the number of cells found together, TNBC cells were further classified in single intravascular (S-IV, white) or in cluster (yellow), particularly, small (2-3 cells, SC-IV) or big (>4cells, BC-IV) intravascular clusters (Scale bars: 50µm) **(C)** Bar plot representation of TBNC cells classification, as a percentage of total cells counted, over time (24, 48 and 72 hours). Data presented as mean ± SD. Statistical analysis was performed using Two-way ANOVA and Sidak’s multiple comparisons test*. P*-value:* <0.05 or “ns” >0.05 (non-significant; N=3).

### 3.5. ERK3 enhances colonization capacity of TNBC cells to lung organoids in *in vitro* models

In the final stages of the metastatic process, the EMT plasticity confers BC cancer cells the ability to colonize the metastatic niche since they retain the ability to undergo the reverse process called mesenchymal-to-epithelial transition (MET)[2]. To further characterize ERK3 role in the acquisition of plasticity by TNBC cells in favour of metastasis, we first investigated its effects in cellular adhesion using decellularized ECM (dECM) from different organs, i.e. lung and liver. Adhesion assay revealed that, overall, cancer cells exhibit greater adhesion to lung compared to liver dECM, consistent with the overall preference for TNBC cells to colonize primarily lung (Fig. 5A).[34,35] Interestingly, while no difference was observed between the two cell types regarding adhesion to liver dECM, shWT cells demonstrated increased adhesion to lung dECM compared to shERK3 cells. These results suggest that ERK3 modulates cell adhesion in an ECM-dependent manner, which differs from tissue-to-tissue.

**Figure 5.**
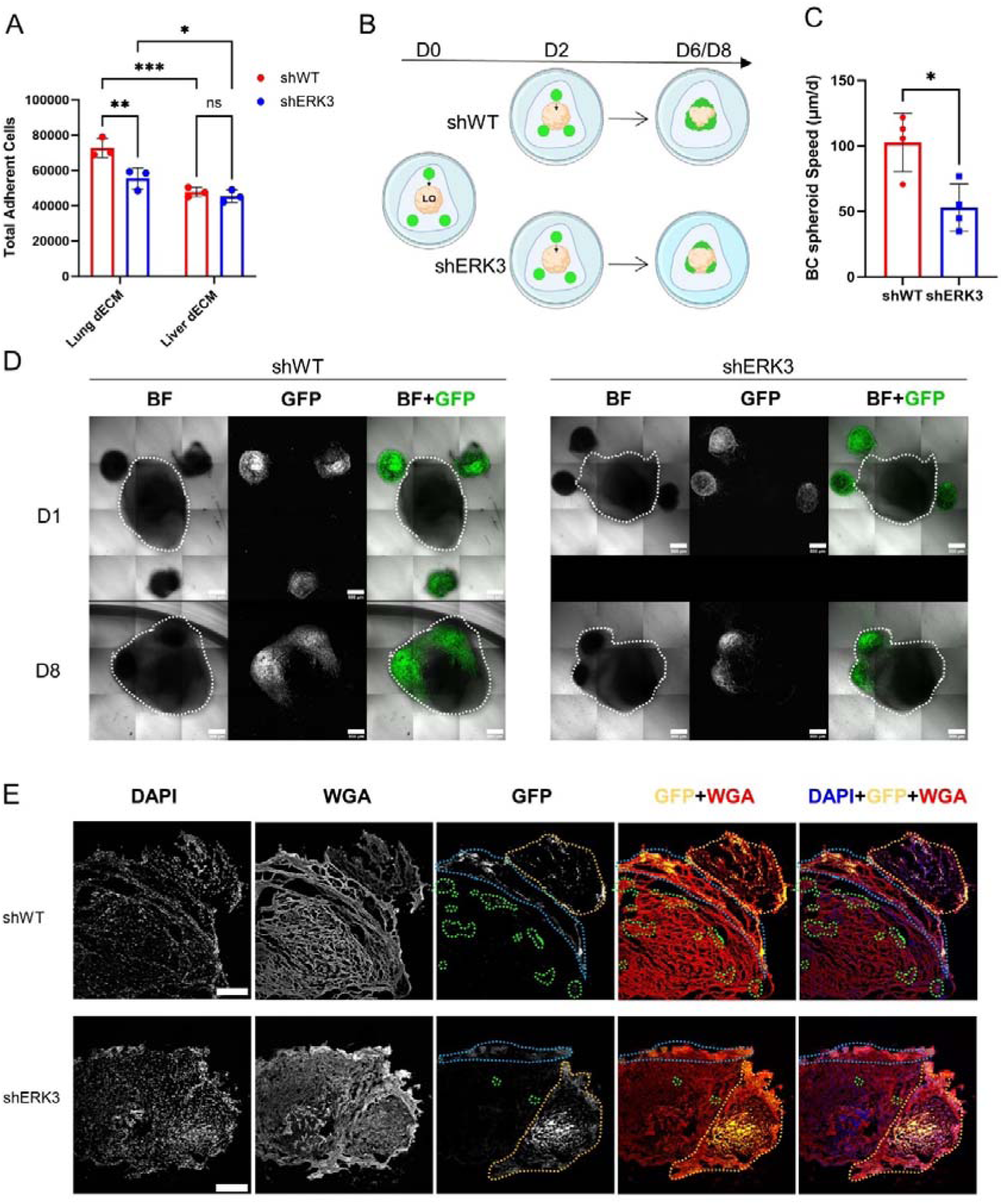
ERK3 promotes TBNC invasion to lung in 3D *in vitro* models. **(A)** TNBC cells adhesion to lung dECM (purple bars) or liver dECM (green bars). **(B)** Schematic of the confrontation assay, in which iPSC-derived lung organoids (LO, beige) were co-cultured in cultrex with the BC spheroids (in green) for a total of 6-8 days. **(C)** Average speed (µm/h) of TNBC spheroids migration towards the LO estimated by the distances measured from confocal images. **(D)** Representative images of the co-cultures obtained by confocal microscopy for shWT (left) and shERK3 (right) spheroids. Bright field (BF) and GFP labelled cells at day 1 (D1) and after 8 days (D8) are shown (Scale bar: 500µm). Merge channel is also shown. **(E)** Representative confocal images of samples obtained from the confrontation assay as in (D), cultured for 8 days. The cryosections were stained with DAPI (blue) and WGA (red). Dashed orange line represents the spheroid, blue line highlights the area of adhesion of the spheroid to the surface of the LO and green dashed line highlights penetrating cells or clusters of cells (Scale bar: 250µm). Data presented as mean ± SD. Statistical analysis was performed using two-way ANOVA followed by Tukey’s multiple comparisons test (panel A, N=3) or unpaired student t-test (panel C, N=4 -shWT n=8; shERK3 n=5). * P <0.05,**P <0.01, ***P <0.001

**Figure 6.**
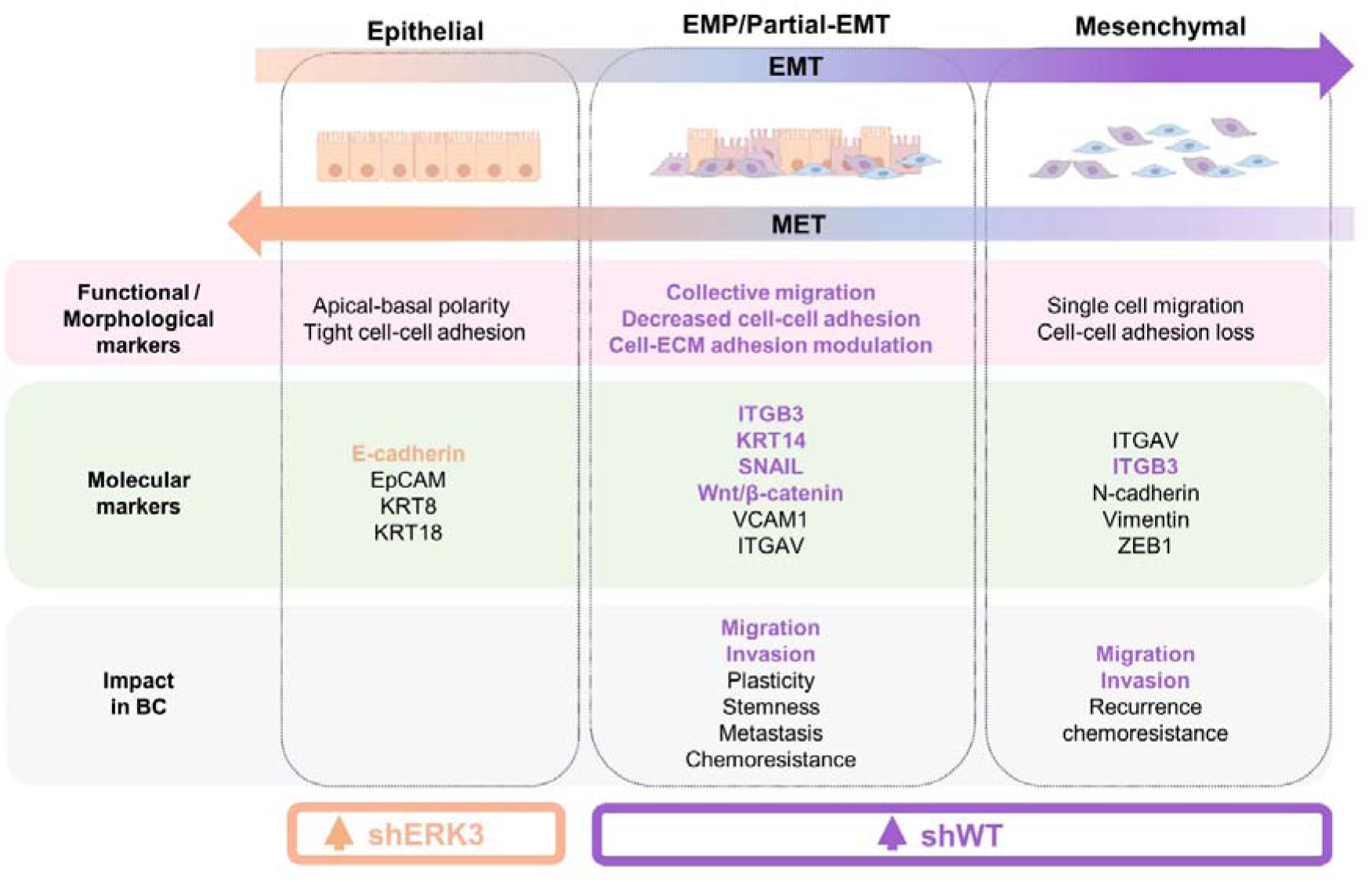
Summary of the different cellular states occurring during EMT based on morphological, functional and molecular markers, and their impact in BC progression. Markers and impact of EMP and full EMT in BC are listed based on recent evidence[3–5,42,43]. Text is highlighted based on the characterization of ERK3 effect on these markers in our shWT (in purple) and in our shERK3 cells (in beige).

In the previous experiments we established that ERK3 participates in promoting migration and invasion in 3D models (Fig. 2F and 3G), and adhesion to lung dECM (Fig. 5A). To deepen our understanding of this process and contextualize within the metastatic process, we investigated the colonization capacity of TNBC spheroids in 3D by co-culturing them with hiPSC-derived lung organoids (LO), and performing a confrontation assay[24]. This methodology has been previously described in the literature to study the migration and invasion capacity of cancer cells[60]. Firstly, the GFP-tagged shWT or shERK3 TNBC spheroids were embedded in cultrex in the proximity of the LO and then the migration of the TNBC spheroids towards the LO was monitored for 6-8 days (Fig. 5B). The approximate speed of the spheroid migration was extrapolated by calculating the distance between the surface of each BC spheroid and the surface of the LO, daily. Of note, TNBC spheroids which were placed at longer distances from the LO (approx. >500 µm) did not migrate (respective spheroids are identified by blue numbers in S4.1A-S4.4A, specifically shWT spheroids 3, 4 and 8; shERK3 spheroids 4 and 9). For all the other spheroids (distance <500 µm, identified by yellow number in S4.1A-S4.4A, specifically shWT spheroids 1, 2, 5, 6, 7, 9, 10 and 11; shERK3 spheroids 3, 5, 7, 10 and 11), the data displayed in Figure 5C shows that shWT spheroids moved faster than shERK3 spheroids (102.715± 22.463 µm/d vs 52.884±18.832 µm/d, respectively). Furthermore, confocal live imaging revealed that, once in contact with the LO, the shWT spheroids adhered to the LO and the cells spread across a larger volume of the LO compared to the cells protruding from shERK3 spheroids (Fig. 5D and S4.1B-S4.4.B). To corroborate this finding, at day 6 or 8 of culture, the co-cultures were fixed, cryosectioned and stained with fluorescent markers for cell membrane (WGA) and the nucleus (DAPI). The results showed that the shWT spheroids infiltrated and colonized deeper into the LO, as depicted by the areas highlighted with the orange, blue and green dashed line, while the cells from shERK3 spheroids remained at the peripheral sections of the LO (Fig. 5E, S4.1C-S4.4.C). Of note, during the placement of the TNBC spheroids, 5 out of 12 shERK3 spheroids were placed in complete contact to the LO (respective spheroids identified by red number in S4.1.A-S4.4.A, specifically shERK3 spheroids 1, 2, 6, 8 and 12), while no shWT spheroid were placed in such proximity. Nonetheless, the shERK3 spheroids presented a significantly lower invasion capacity in comparison to the shWT spheroids (Fig. S4.1.-S4.4.).

These results highlight the interplay between ERK3 and the ECM in modulating cell adhesion, suggesting a reciprocal influence, and show that ERK3 enables efficient collective migration and invasion of TNBC cells towards healthy lung organoids, a likely metastatic site for TNBC cells. Altogether, these results support our hypothesis that ERK3 promotes higher TNBC cell plasticity and adhesion to secondary tissues, i.e. lung.

## 4. Discussion

TNBC is the most aggressive subtype of BC and is characterized by higher cell plasticity, which confers recurrence and metastatic rate, and is evaluated with high tumor grading[34]. In this context, ERK3 has been shown to promote cancer cell migration and metastasis using *in vivo* models[12–14]. However, the specific role of ERK3 during cell migration and colonization of secondary organs in TNBC remains unresolved.

Through the analysis of large multi-omics datasets, we found that ERK3 has a higher gene expression in normal breast tissue compared to the other MAPKs. Typically, ERK3 shows strong expression in glandular epithelia in breast samples[61]. This might be due to ERK3 high turnover and its key role in establishing epithelial architecture in mammary tissue, by modulating MET and cell-cell adhesion[10,62]. Furthermore, we showed that ERK3 is overexpressed in BC patient samples, in both primary tumor and metastatic biopsies. Importantly, this increased expression correlates with aggressive subtypes of BC, such as TNBC, higher tumor grading and worse survival metrics (OS, RFS, DMFS). Therefore, we hypothesized that ERK3 may contribute to increase epithelial-mesenchymal plasticity (EMP) and, consequently, to the metastatic process occurring in TNBC.

Using MDA-MB231 as a model for TBNC, we observed that ERK3 promoted collective migration in a 2D model but had no influence on single-cell migration. Previous studies have reported a pro-migratory effect of ERK3 in lung cancer using the transwell assay[63,64]. Similar findings were reported in a study using BC cells, where EGF was used as a chemoattractant[13]. Here, we used 10% FBS media, instead of a specific chemoattractant, suggesting that ERK3 preferably promotes collective migration of TNBC cells rather than single-cell migration. Interestingly, collective migration is defined as a key feature of a partial-EMT state – or EM plasticity[2–4,43]. Since EMT refers to a spectrum of phenotypes and not to a binary switch, cells with higher plasticity often display intermediate stages of EMT which should be defined by both changes in cellular behaviour and molecular markers[3]. Thus, we investigated additional functional changes in cell behaviour, characteristic of EM plasticity, and observed that ERK3 decreases cell adhesion to different ECM substrates, i.e. collagen IV, laminin, collagen I, fibronectin, tenascin and vitronectin, while promoting anchorage-independent survival. Our results are consistent with previous studies which have demonstrated this pro-survival mechanism in several types of cancer, including breast[14,65].

Next, we investigated the expression of key molecular markers of EMP in TNBC. This intermediate state of EMT is driven mostly by SNAIL transcription factor and canonical Wnt/β-catenin signaling[42]. Additionally, KRT14 is known to be overexpressed in leader cells in collective migration of TNBC, and is also recognized as a marker of EMP [5,43]. Consistently, EMP leads to higher migration of invasive tumor cells through the stroma of the breast tissue[2,5,42,43]. In our study, we demonstrate that ERK3 knockdown reduces KRT14 protein expression and downregulates, at both transcriptional and protein level, SNAIL and β-catenin in TNBC cells. Additionally, we demonstrate that the loss of ERK3 also downregulates at both protein and transcriptional level YAP, which has been shown to interact with β-catenin in KRT14+ cells, inducing and sustaining CSC survival, EMT and overall TNBC progression[66]. Moreover, we observed a reduction in the expression of the E-cadherin gene (CDH1), which is typically repressed by SNAIL, in our control cells (shWT) compared to ERK3 knockdown cells (shERK3). We also noted the downregulation of downstream effectors of β-catenin signaling, WISP1 (CCN4), and of YAP, such as CYR61 (CCN1) and CTGF (CCN2), which are implicated in inducing EMT[67,68].

Once established the contribution of ERK3 to enhance EMT plasticity through molecular and functional characterizations, we inquired whether this mechanism contributes to higher metastatic capacity in TNBC cells. By using different advanced *in vitro* models, including organoids and microfluidic systems, we aimed to mimic the lung tissue – the preferred metastatic site of TNBC – and study how cancer cells migrate towards the healthy lung tissue model and colonize it[34,35]. Firstly, we investigated the extravasation capacity using a 3D-MPS that mimics human lung capillaries. While we observed lower extravasation rate compared to a previously reported study, which we attribute to the absence of constant flow in our system, our study revealed that ERK3 increases mid-extravasation of cancer cells after 72 hours[30]. As mentioned before, ERK3 increases both at protein and transcriptional level CYR61, which has been demonstrated to increase BC extravasation, possibly by promoting survival of circulating tumor cells[69]. In particular, CYR61 is overexpressed in relapsed TNBC tumors and metastatic biopsies[70]. We also observed increased mRNA expression of one of CYR61s receptors – ITGB3/CD61 – which is a marker for late EMT stage and CSC[43,70]. Secondly, using decellularized ECM of either lung or liver, we have demonstrated that ERK3 preferably promotes cellular adhesion to lung dECM, highlighting the interplay between ERK3-induced EMT plasticity and ECM specificity. Finally, we co-cultured human iPSC-derived lung organoids with TNBC cells, establishing an *in vitro* model of organ colonization. This methodology has been previously developed by our group to study the metastatic potential of patient-derived prostate cancer organoids towards lung organoids[60]. Herein, we used GFP-labelled tumor cells for tracking spheroid migration towards the LO as well as to evaluate TNBC cells penetration capacity. Our observations showed the TNBC spheroids migrating cohesively toward the LO, rather than radially, as described for the 3D invasion assay in collagen I and cultrex. Interestingly, using confocal microscopy and ImageJ to analyse the results from the confrontation assay, we demonstrated that ERK3 promotes faster spheroid migration towards the LO, as quantified by approximately 2-fold faster average speed (µm/d) for shWT in comparison with shERK3 cells. Additionally, through a qualitative approach, we observed deeper penetration into LO of TNBC cells expressing ERK3 compared to those where ERK3 was knocked down. These findings support the use of this model for future studies of cancer cell colonization and importantly corroborates our hypothesis that ERK3 pro-migratory and pro-invasive role contributes to metastasis of TNBC, as previously reported in *in vivo* models[12–14].

In conclusion, our study shows that ERK3 promotes epithelial-mesenchymal plasticity in TNBC through the deregulation of several key molecular markers, including KRT14 and SNAIL, and important signaling pathways such as β-catenin, YAP and CYR61. These alterations are supported by functional changes towards partial-EMT state characterized by the modulation of cell-ECM adhesion, anchorage-independent survival, collective migration and invasion. Using advanced 3D *in vitro* models, we corroborated the pro-metastatic behaviour of ERK3 in TNBC cells, particularly towards lung colonization. The importance of our study is highlighted by the overexpression of ERK3 in more plastic and aggressive patient tumor samples and by its contribution to poor patient survival. Here we were able to contribute to the understanding of how ERK3 promotes TNBC progression and possible metastization, warranting future studies to elucidate if ERK3 could be a valuable therapeutic target or biomarker in breast cancer.

## Abbreviations

(EMP): Epithelial-mesenchymal plasticity
(3D-MPS): 3-dimensional microphysiological system

## Data availability

All data generated during this study are available upon request.

## Authors contributions

S.M. and G.F. conceptualized the study. S.M, S.F. and G.F. wrote the manuscript. S.M. designed the methodology, performed data analysis and data curation. S.F., M.F., H.S., M.C., K.V, J.V., DPS, V.B. and J.S. assisted in methodology and data analysis. M.F. and K.H. provided the methodology and supervision to generate the 3D microfluidic systems for the extravasation assay. F.K., V.B. and J.F. provided the methodology and supervision to generate and characterize the hiPSCs-derived lung organoids. All authors contributed to edit and review the manuscript, and approved the final version. G.F. supervised this study.

## Acknowledgements

This work was supported by Ministry of Health of the Czech Republic, grant no. NU23J-08-00035, by Marie Curie H2020-MSCA-IF-2020 MSCA-IF-GF “MecHA-Nano”, Grant Agreement No 101031744 and DRO (Institute of Hematology and Blood Transfusion – UHKT, No. 00023736). Moreover, this work was also supported by the European Regional Development Fund-Project MAGNET (Number CZ.02.1.01/0.0/0.0/15_003/ 0000492) and ENOCH (No. CZ.02.1.01/0.0/0.0/16_019/0000868). This work received funding from the European Union’s Horizon 2020 research and innovation programme under grant agreement No. 860715 and No.101104587. Additionally, it was also supported by EMBL internal funds. Sofia Morazzo was supported by the EMBL Corporate Partnership Programme fellowship for scientific visitors and by Scholarships of the Ministry of Education, Youth and Sports in Support of Foreign Nationals’ Study at Public Institutions of Higher Education in the Czech Republic (MSMT-9985/2021-4-001 and MSMT-8872/2022-1-001). Giancarlo Forte was supported by the King’s BHF Centre for Excellence Award (RE/18/2/34213).

We are thankful to to Jaromíra Novaková for her help with cell culture and western blots, Jirí Smíd for his help with sample cryosectioning, and Romana Vlčková and Hana Dulová for the continuous technical support. Illustrations were created with BioRender (https://www.biorender.com/).

---

**Supplementary Figure 1.**
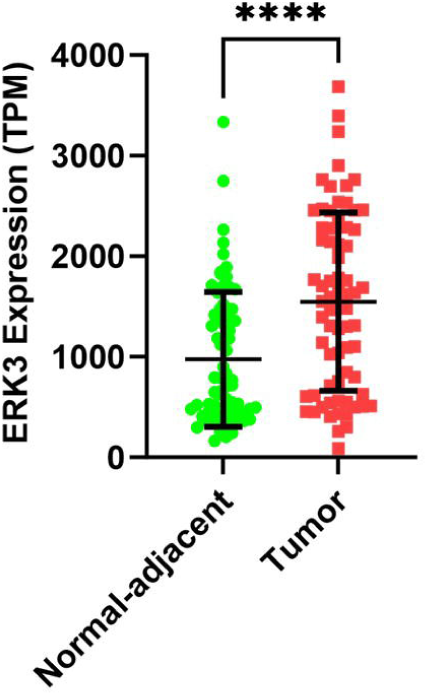

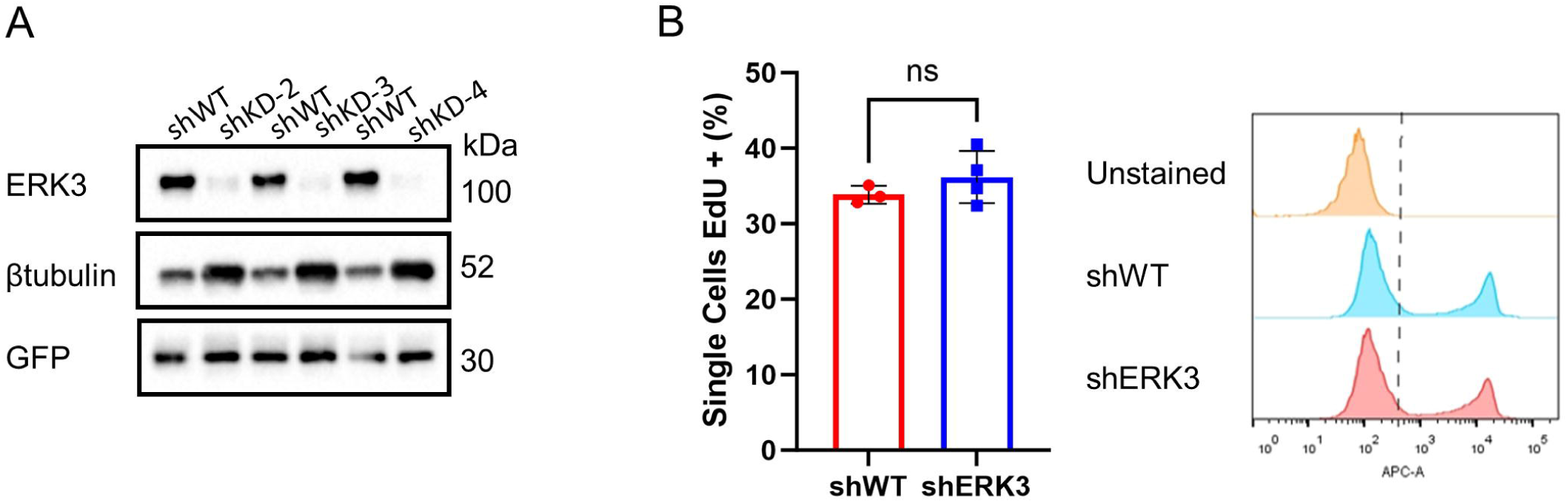

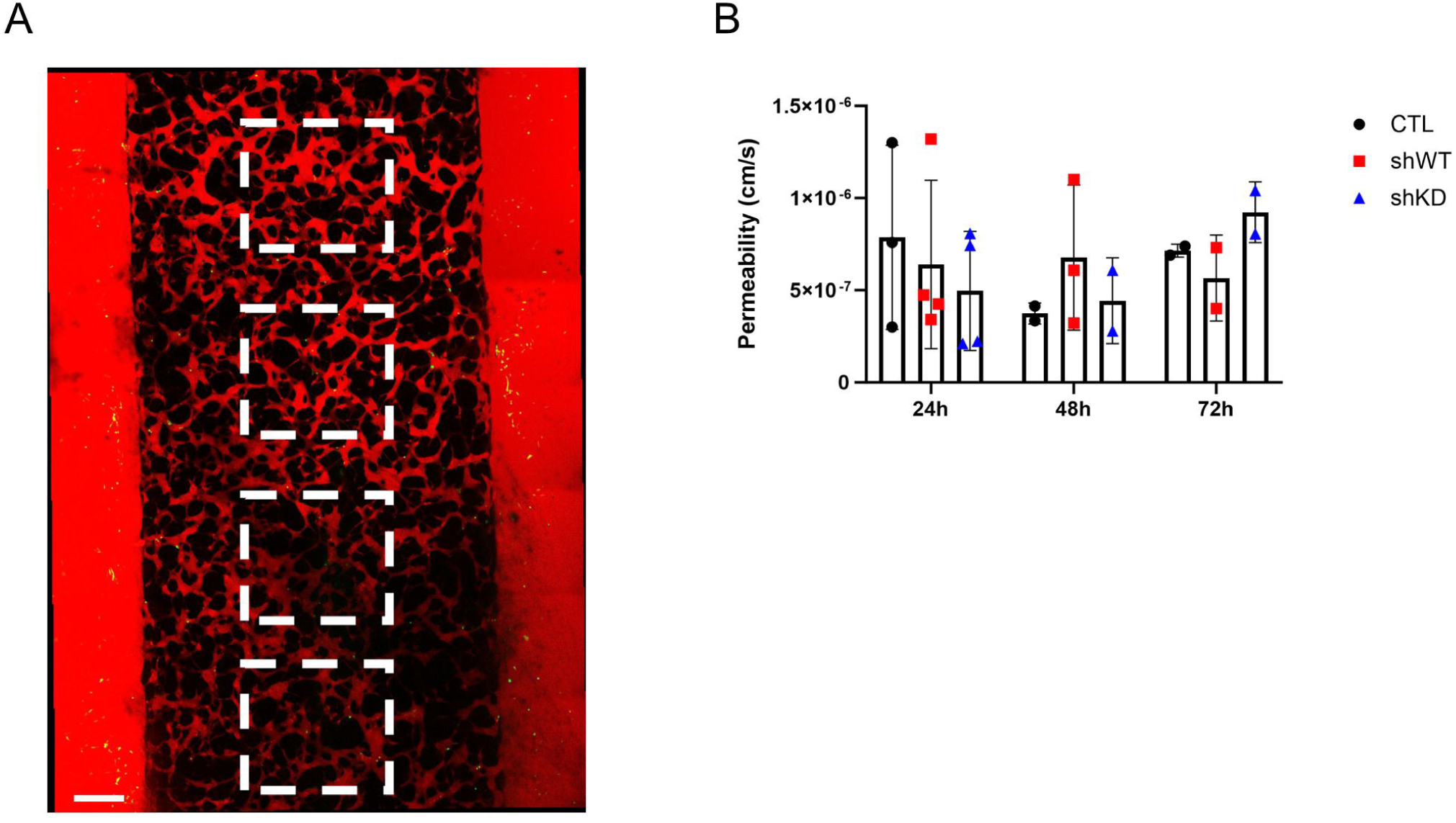

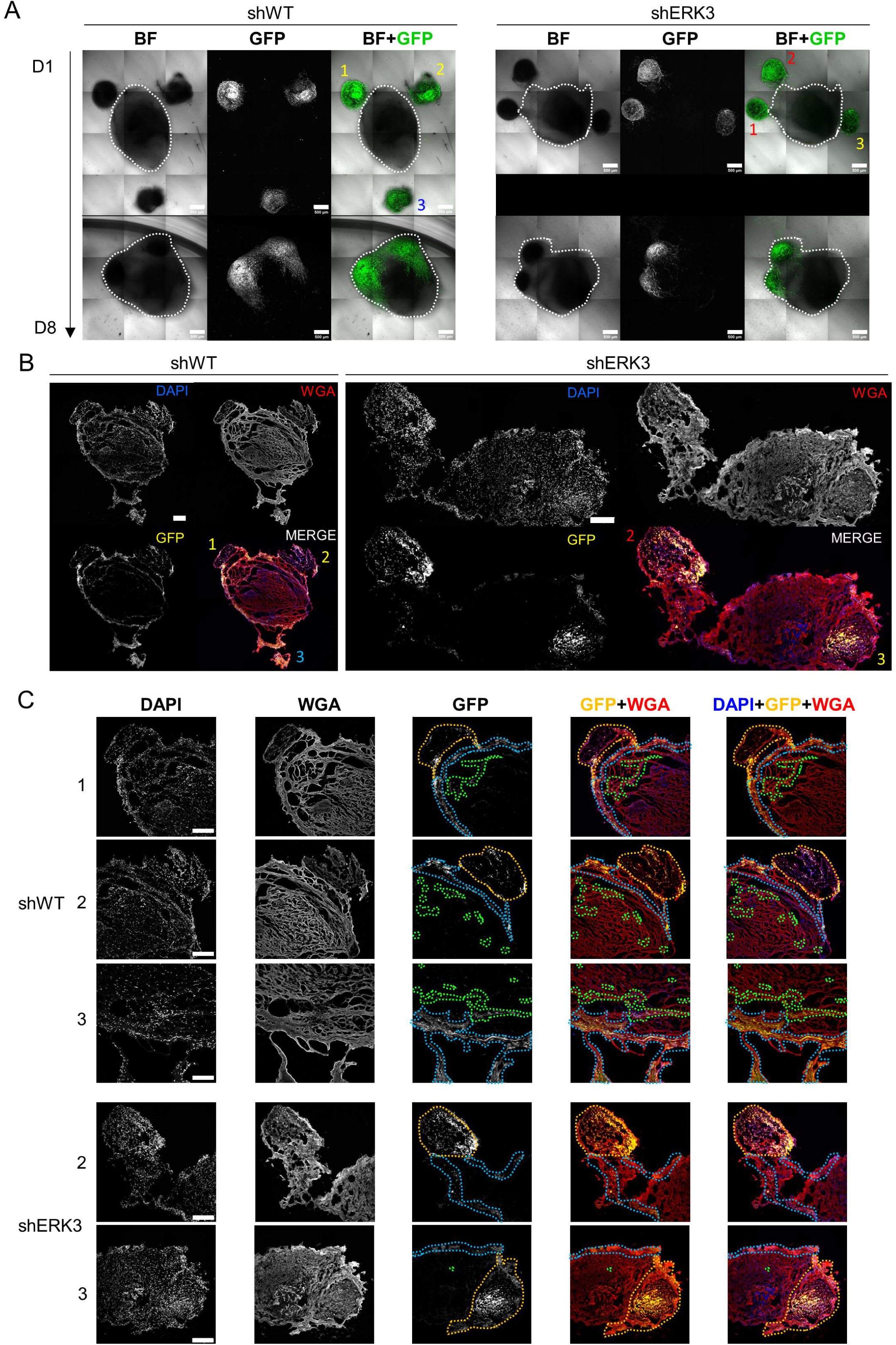

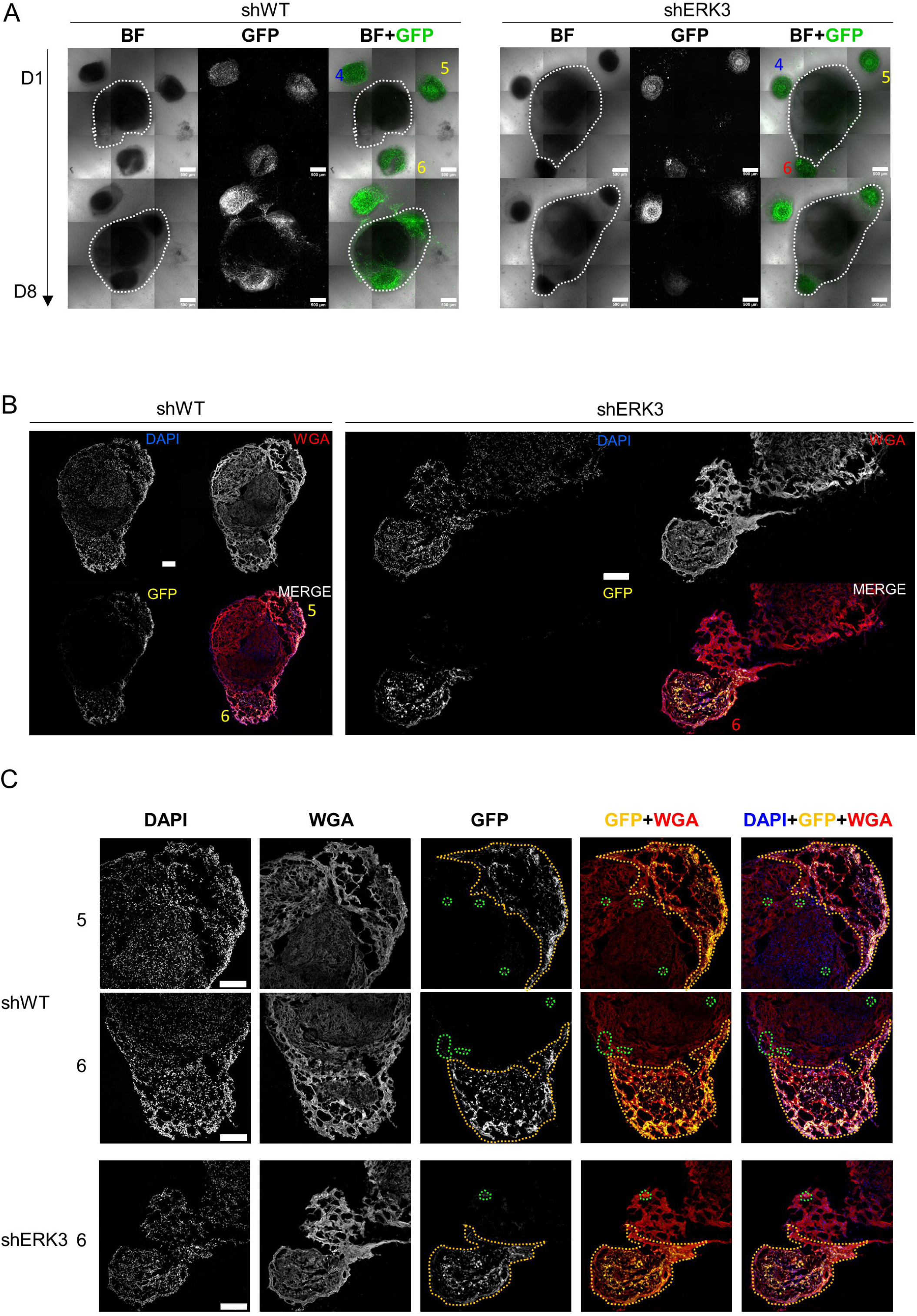

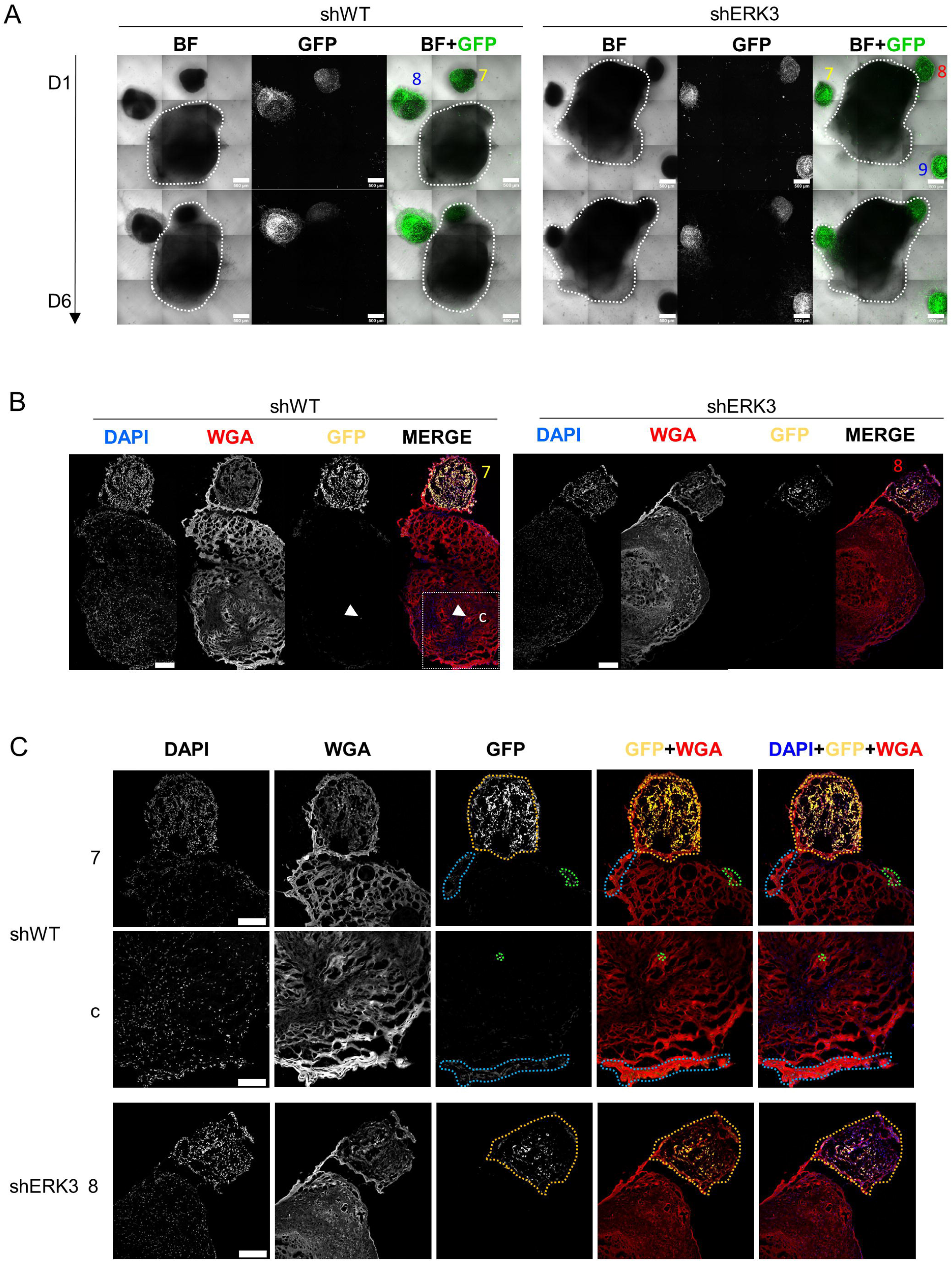

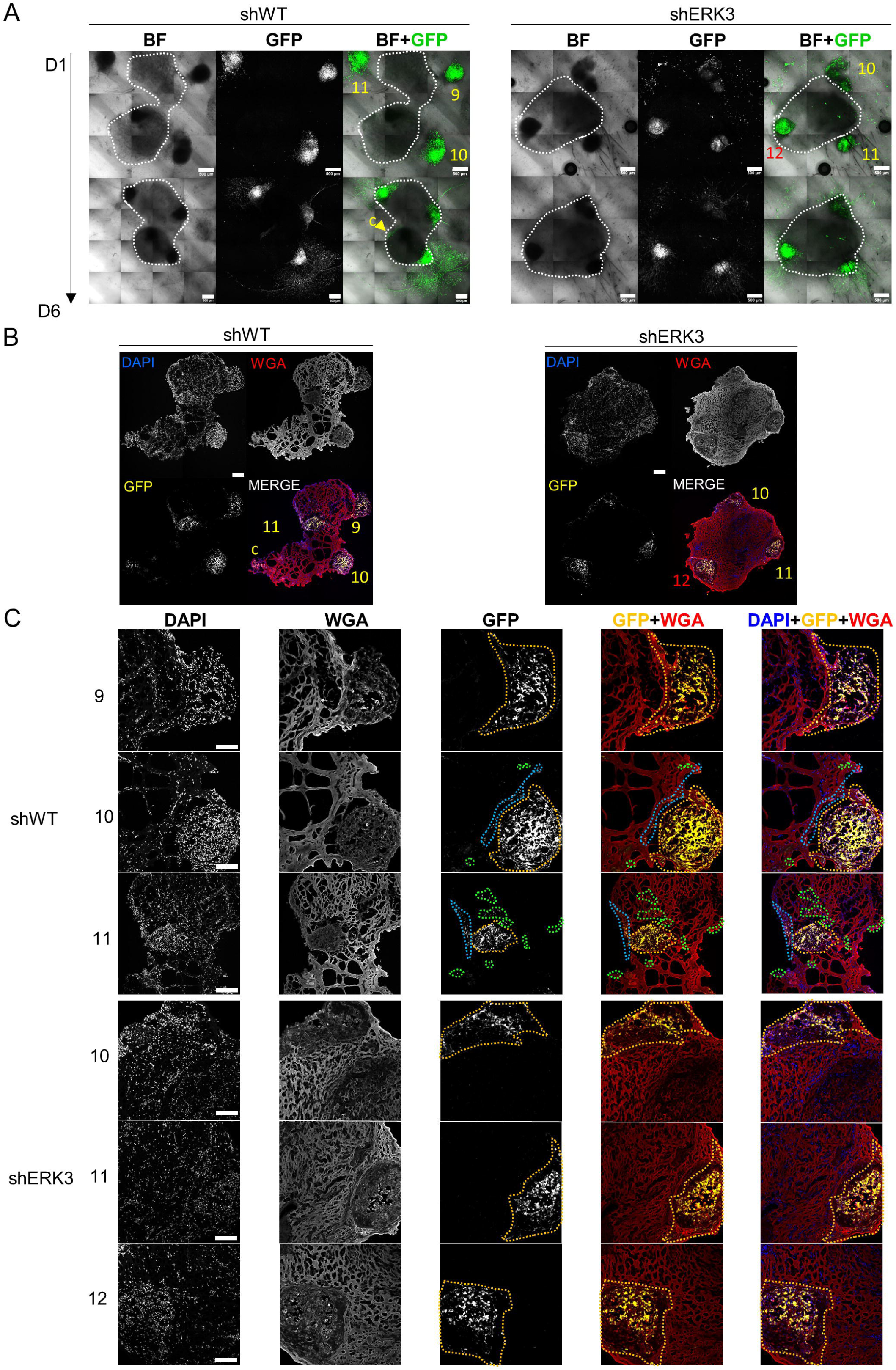

