## Supplementary Material for "ERK3/MAPK6 promotes triple-negative breast cancer progression through collective migration and EMT plasticity"

### Abstract

Triple-negative breast cancer (TNBC) is the most aggressive subtype of breast cancer and is associated with high cell plasticity, recurrence, and metastatic rate. During epithelial-to-mesenchymal transition (EMT), cancer cells display EMT plasticity, or partial-EMT features, which are required for breast cancer metastasis, such as collective migration. ERK3 has been implicated in promoting migration and invasion of breast cancer, but the mechanisms remain elusive. Here, we investigated ERK3 expression across patient-derived datasets of breast cancer and established its association with aggressive breast cancer phenotypes and poor clinical outcomes. Leveraging the hypothesis that ERK3 contributes to TNBC progression by supporting a partial-EMT state, we showed that ERK3 is essential in different steps of the metastatic process, especially by enabling collective migration but also by modulating cell-extracellular matrix adhesion, anchorage-independent growth, extravasation and colonization. In conclusion, our results demonstrate that ERK3 contributes to TNBC progression and potentially metastasis by promoting EMT plasticity and collective migration.

**Keywords:** triple-negative breast cancer (TNBC), extracellular-regulated kinase 3 (ERK3), epithelial-to-mesenchymal transition (EMT), epithelial–mesenchymal plasticity (EMP), collective migration.

### Abbreviations

Epithelial-mesenchymal plasticity (EMP)

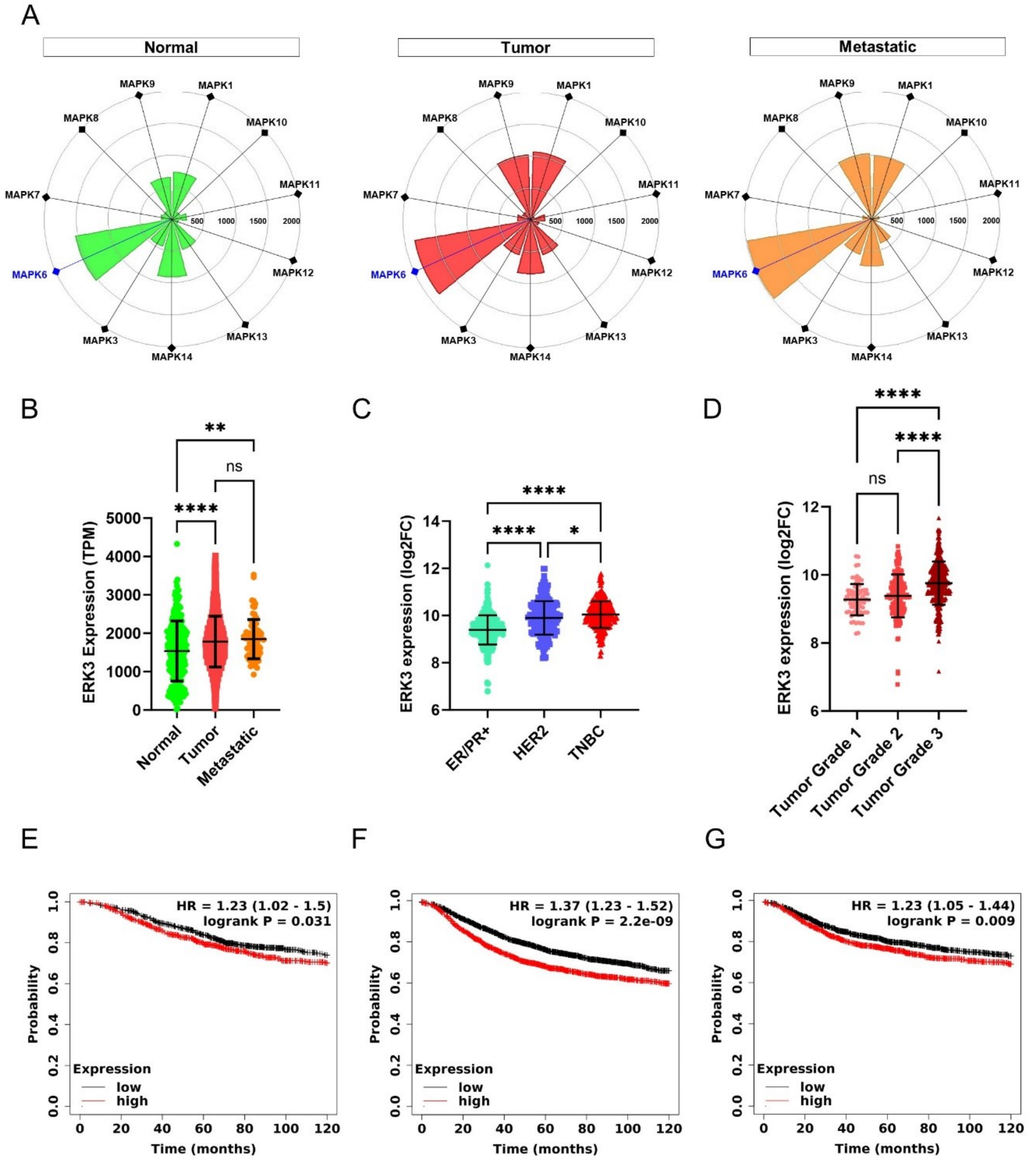


**Figure 1. ERK3 overexpression in BC patients correlates with poor patient survival.** (**A**) Targetgram analysis of the average expression (transcripts per million, TPM) of the different MAPK genes, in normal breast tissue and in primary and metastatic breast tumors, extracted directly from TNMplot[21]. **(B)** Dotplot representation of ERK3 gene expression (TPM), in normal breast tissue and in primary and metastatic breast tumors (Normal-N=242, Tumor- N=7440, Metastatic- N=77), from the TNMplot. ERK3 gene expression is shown as log2 fold change (log2FC), from the GENT2 platform **(C)** distributed by molecular subtype (ER/PR+ - N=623, HER2 – N=229, TNBC – N=251), and (**D**) by BC histological grade (grade 1 – N=63, grade 2 – N=151, grade 3 – N=358). Survival curves showing the probability of overall survival – (OS)**(E)**, recurrence-free survival (RFS)**(F)** and distant-metastasis free survival (DMFS)**(G)** of BC patients with either high (red) or low (black) expression of ERK3. Accessed and analyzed through the Kaplan-Meier plotter platform. Dotplot graphs show data as mean ± SD. Statistical analysis was performed either by one-way ANOVA followed by Tukey's multiple

comparisons test (B, D and E), paired student t-test (C) or by log-rank test (F, G and H). HR – hazard ratio. * $P < 0.05$, **$P$<0.01, ****$P$<0.0001 or ns – non-significant.

Altogether, these results show that ERK3 promotes EMP features associated with metastasis by supporting collective migration and enhancing survival and invasion capabilities of TNBC cells.

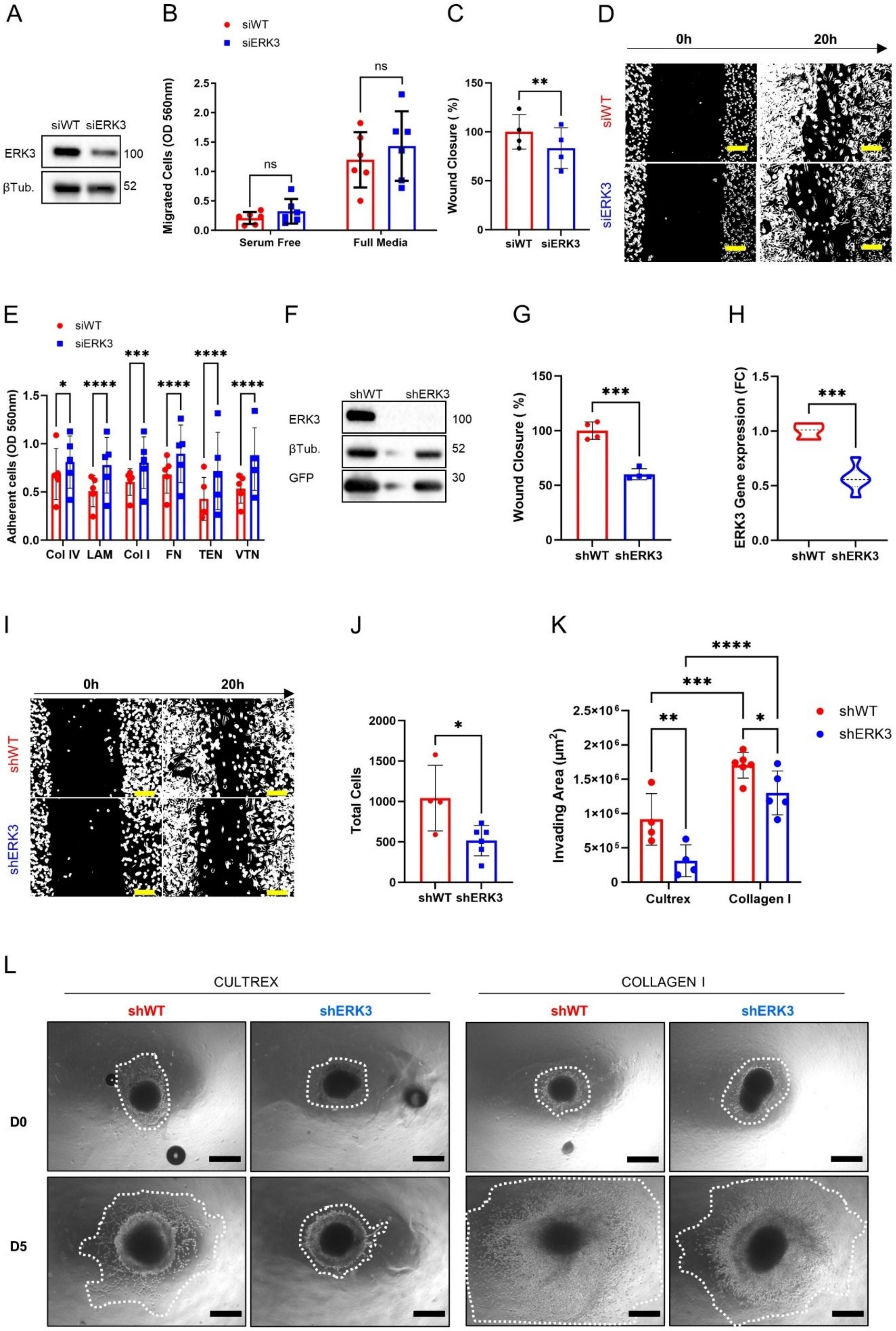


**Figure 2 – ERK3 promotes collective migration and invasion of TNBC cells. (A)** Transient ERK3 knockdown in MDA-MB231 cell line by siRNA and respective scramble control (siERK3 and siWT, respectively) was validated by western blot, for ERK3 and β-tubulin (βTub.). **(B)** Transwell migration assay using siERK3 and siWT cells in either serum-free or full media conditions (N=6 for 20h). **(C)** wound healing assay using siERK3 and siWT cells (N=4) and **(D)** respective representative image of the wound healing assay at 0 h and 20 h (Scale bar: 100µm). **(E)** siERK3 and siWT cells-ECM adhesion assay, across different ECM substrates (Col IV – collagen type IV, LAM – laminin, Col I – collagen type I, FN – fibronectin, TEN – tenascin, VTN – vitronectin; N=5). **(F)** and **(G)** Stable MDA-MB231 cell lines with and without shRNA-induced ERK3 silencing (shERK3 and shWT, respectively), were validated by western blot for ERK3, β-tubulin (βTub.) and GFP **(F)**, and by qPCR for ERK3 gene expression (N=3) **(G)**. **(H)** and **(I)** Wound healing assay performed using shERK3 and shWT stable cell lines (N=4), as characterized by qPCR analysis **(H)**, and representative image of the wound healing assay at 0 h and 20 h **(I)** (Scale bar: 100µm). **(J)** Total cell number determined in soft-agar colony formation assay, after 8 days in culture (shWT - N=4, shERK3 – N=6). **(K)** and **(L)** Migration and invasion ability determined by quantifying the invading area of TNBC spheroids at day 0 and day 5 of culture, in either cultrex or collagen (cultrex - N=4; collagen I – N=6) (K), and representative images with the corresponding invading area **(L)** (white dotted lines; Scale bar: 250µm). All data are presented as mean ± SD. Statistical analysis was performed either by paired student t-test (B and C), unpaired student t-test (F, G and I), or two-way ANOVA followed by by Šídák's multiple comparisons test (D and J) * $P < 0.05$, ** $P < 0.01$, *** $P < 0.001$, **** $P < 0.0001$ or ns – non-significant .

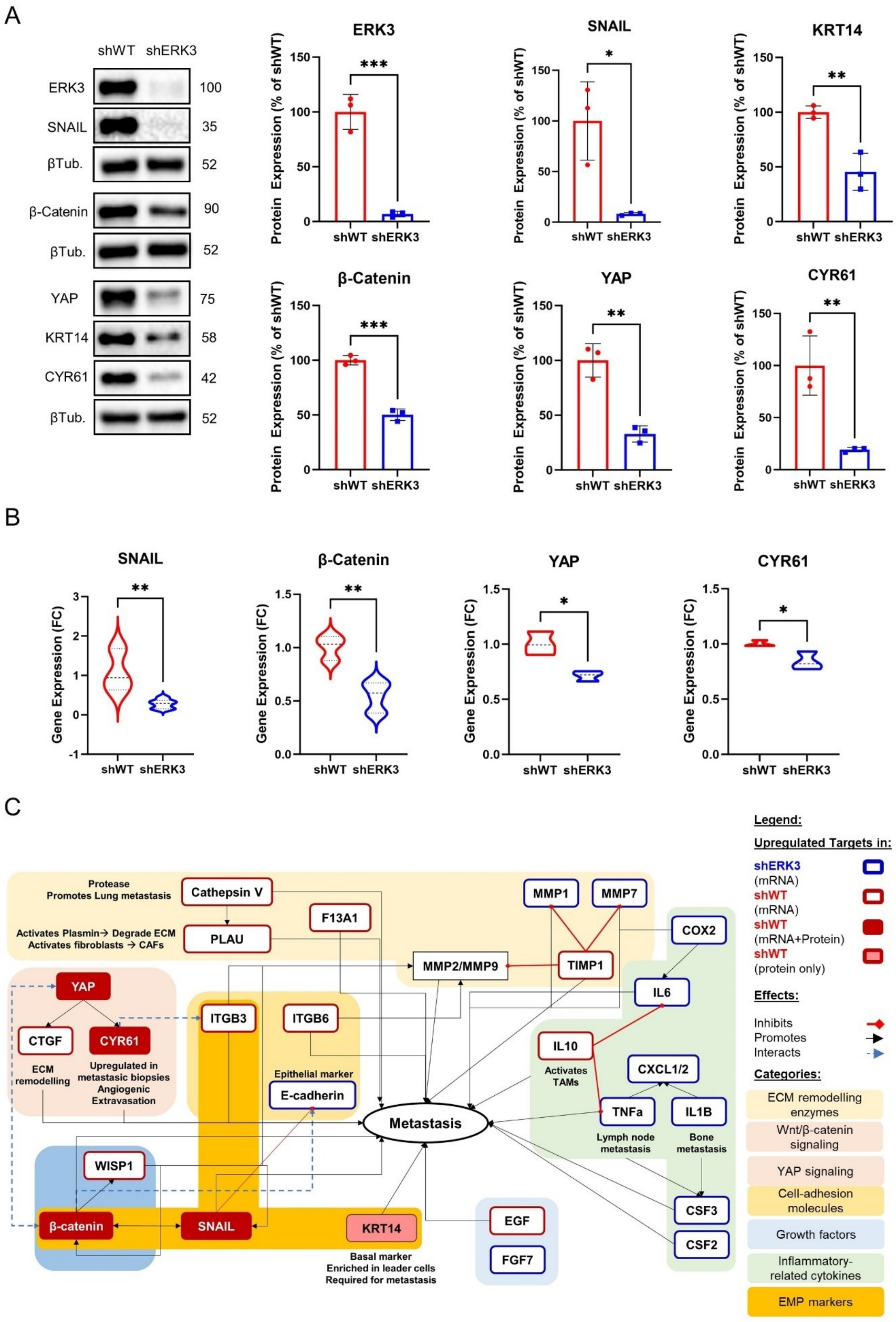
A
shWT shERK3
ERK3 100
SNAIL 35
βTub. 52
β-Catenin 90
βTub. 52
YAP 75
KRT14 58
CYR61 42
βTub. 52
ERK3
SNAIL
KRT14
β-Catenin
YAP
CYR61
Protein Expression (% of shWT)
shWT shERK3
***
*
**
B
SNAIL
β-Catenin
YAP
CYR61
Gene Expression (FC)
shWT shERK3
C
Protease
Promotes Lung metastasis
Cathepsin V
Activates Plasmin→ Degrade ECM
Activates fibroblasts → CAFs
PLAU
F13A1
MMP1
MMP7
COX2
MMP2/MMP9
TIMP1
IL6
YAP
CTGF
CYR61
ECM remodelling
Upregulated in metastasic biopsies
Angiogenic
Extravasation
ITGB3
ITGB6
Epithelial marker
E-cadherin
IL10
Activates TAMs
CXCL1/2
TNFa
IL1B
Lymph node metastasis
Bone metastasis
Metastasis
WISP1
β-catenin
SNAIL
KRT14
Basal marker
Enriched in leader cells
Required for metastasis
EGF
FGF7
CSF3
CSF2
Legend:
Upregulated Targets in:
shERK3 (mRNA)
shWT (mRNA)
shWT (mRNA+Protein)
shWT (protein only)
Effects:
Inhibits
Promotes
Interacts
Categories:
ECM remodelling enzymes
Wnt/β-catenin signaling
YAP signaling
Cell-adhesion molecules
Growth factors
Inflammatory-related cytokines
EMP markers

**Figure 3 – ERK3 upregulates EMT, pro-metastatic and collective migration markers in TNBC cells. (A)** Representative images of western blot and relative protein quantification of ERK3, SNAIL, β-catenin, YAP, CYR61, cytokeratin 14 (KRT14), and β-tubulin (βTub.) in shERK3 and shWT stable cell lines. **(B)** Gene expression determined by qPCR and expressed as fold change (FC) of SNAIL, β-catenin, YAP and CYR61 in shERK3 and shWT stable cell lines **(C)** Schematic representation of the differentially regulated markers by ERK3 as determined by $RT^2$ profiler PCR Array, western blot and qPCR, and how they relate to the EMT and/or metastatic process in BC, as reported in literature. Data presented mean ± SD. Statistical analysis was performed using unpaired *t-test.* (N=3). * $P$ <0.05, ** $P$ <0.01, *** $P$ <0.001

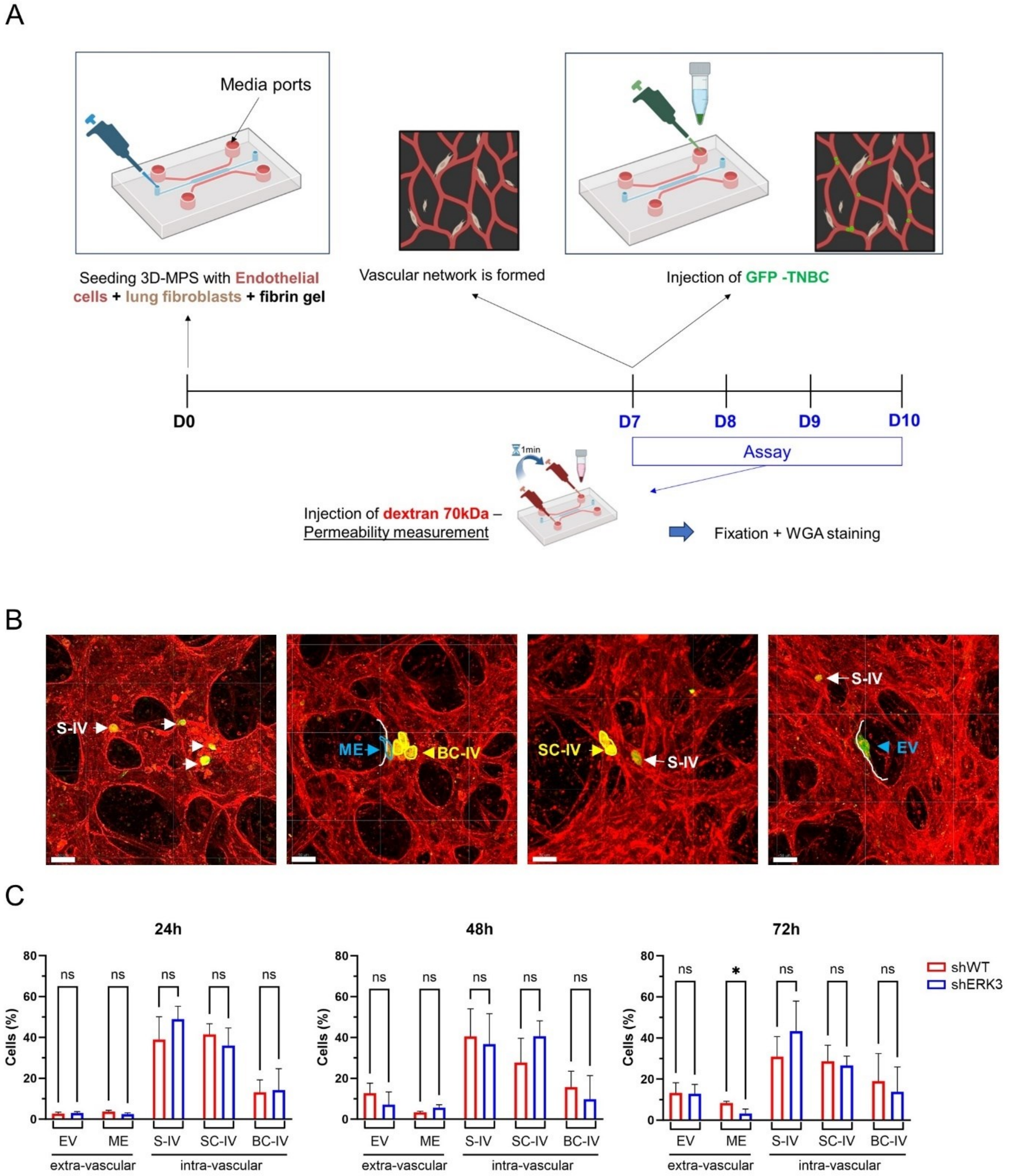


**Figure 4 – ERK3 increases cancer cell extravasation in a 3D microphysiological system, which mimics lung capillaries. (A)** Schematic representation of the 3D-MPS with a central channel (blue) used for the seeding of endothelial cells (red) and lung fibroblasts (brown) within a fibrin gel (top). The parallel media ports used for the injection are indicated by an arrow. Experimental timeline representation of the assay is also shown (bottom). **(B)** Representative confocal images of vasculature stained with WGA (red). GFP-TNBC cells are in green. Depending on their position relative to the blood vessel wall (white line), TNBC cells were classified in intravascular (IV) extravascular (EV) or mid-extravasation (ME, blue). Additionally, depending on the number of cells found together, TNBC cells were further classified in single intravascular (S-IV, white) or in cluster (yellow), particularly, small (2-3 cells, SC-IV) or big (>4cells, BC-IV) intravascular clusters (Scale bars: 50μm) **(C)** Bar plot representation of TBNC cells classification, as a percentage of total cells counted, over time (24, 48 and 72 hours). Data presented as mean ± SD. Statistical analysis was performed using Two-way ANOVA and Sidak's multiple comparisons test. *P*-value:* <0.05 or "ns" >0.05 (non-significant; N=3).

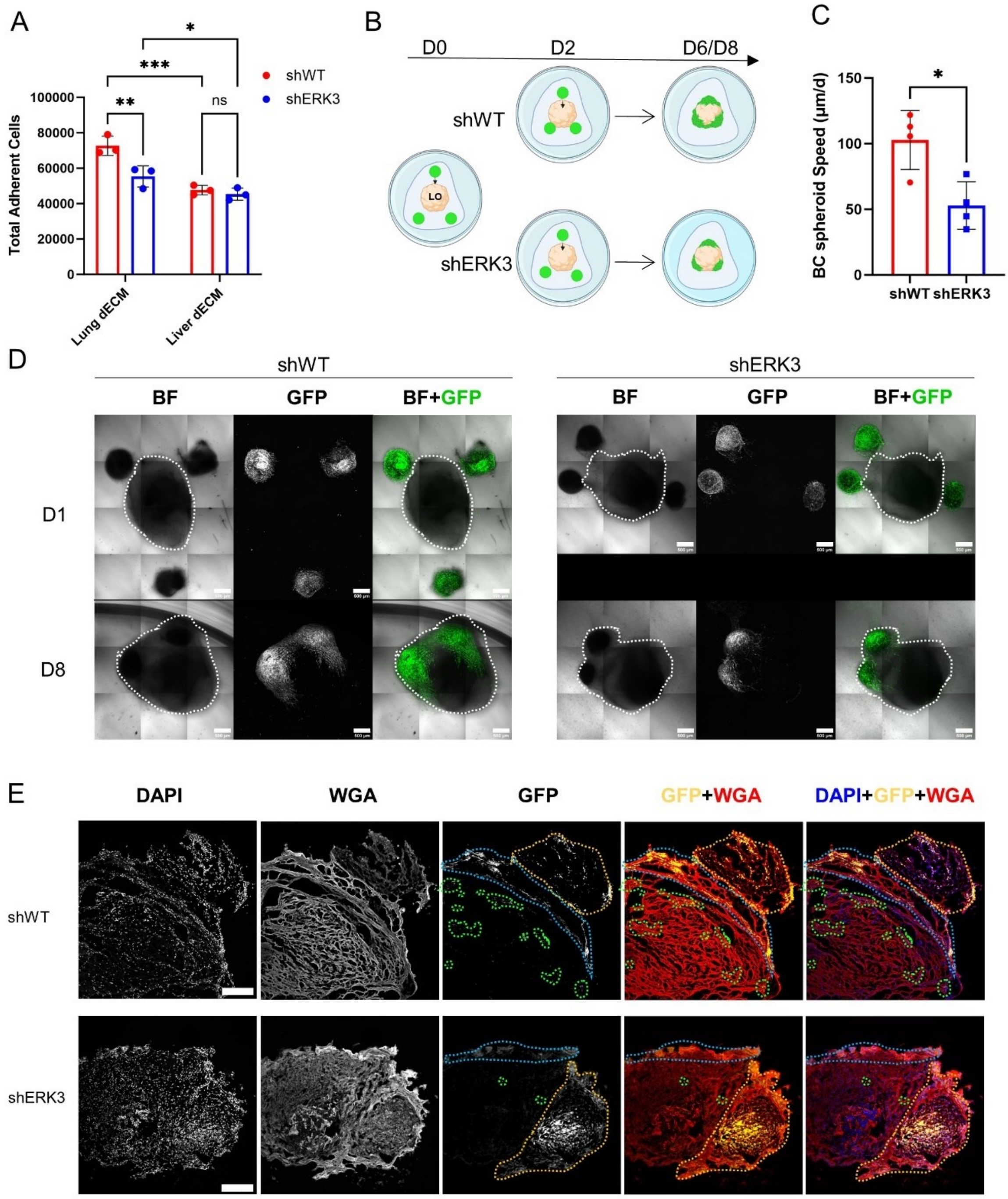


**Figure 5. ERK3 promotes TBNC invasion to lung in 3D *in vitro* models**. **(A)** TNBC cells adhesion to lung dECM (purple bars) or liver dECM (green bars). **(B)** Schematic of the confrontation assay, in which iPSC-derived lung organoids (LO, beige) were co-cultured in cultrex with the BC spheroids (in green) for a total of 6-8 days. **(C)** Average speed (µm/h) of TNBC spheroids migration towards the LO estimated by the distances measured from confocal images. **(D)** Representative images of the co-cultures obtained by confocal microscopy for shWT (left) and shERK3 (right) spheroids. Bright field (BF) and GFP labelled cells at day 1 (D1) and after 8 days (D8) are shown (Scale bar: 500µm). Merge channel is also shown. **(E)** Representative confocal images of samples obtained from the confrontation assay as in (D), cultured for 8 days. The cryosections were stained with DAPI (blue) and WGA (red). Dashed orange line represents the spheroid, blue line highlights the area of adhesion of the spheroid to the surface of the LO and green dashed line highlights penetrating cells or clusters of cells (Scale bar: 250µm). Data presented as mean ± SD. Statistical analysis was performed using two-way ANOVA followed by Tukey's

multiple comparisons test (panel A, N=3) or unpaired student t-test (panel C, N=4 - shWT n=8; shERK3 n=5). * P <0.05,**P <0.01, ***P <0.001

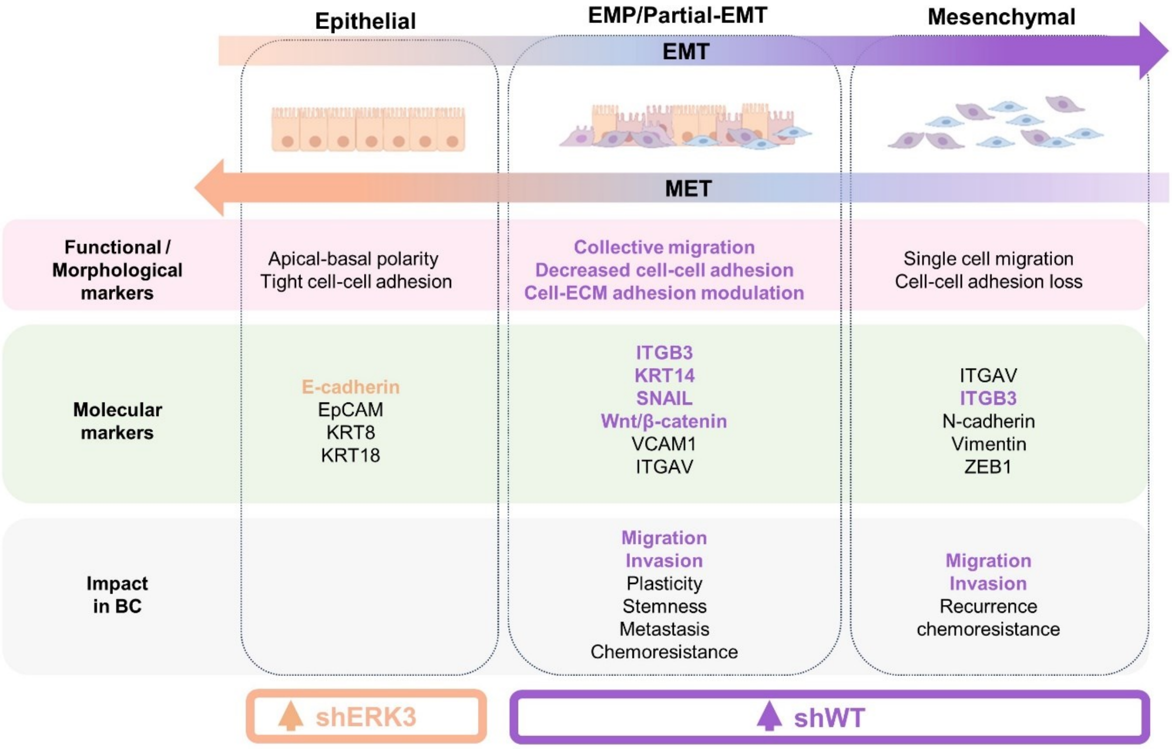


**Figure 6. Summary of the different cellular states occurring during EMT based on morphological, functional and molecular markers, and their impact in BC progression.** Markers and impact of EMP and full EMT in BC are listed based on recent evidence[3–5,42,43]. Text is highlighted based on the characterization of ERK3 effect on these markers in our shWT (in purple) and in our shERK3 cells (in beige).

#### Disclosure of conflicts of interest

The authors declare no conflict of interests.

[18] Lonsdale J, Thomas J, Salvatore M, Phillips R, Lo E, Shad S, Hasz R, Walters G, Garcia F, Young N, Foster B, Moser M, Karasik E, Gillard B, Ramsey K, Sullivan S, Bridge J, Magazine H, Syron J, Fleming J, Siminoff L, Traino H, Mosavel M, Barker L, Jewell S, Rohrer D, Maxim D, Filkins D, Harbach P, Cortadillo E, Berghuis B, Turner L, Hudson E, Feenstra K, Sobin L, Robb J, Branton P, Korzeniewski G, Shive C, Tabor D, Qi L, Groch K, Nampally S, Buia S, Zimmerman A, Smith A,

Burges R, Robinson K, Valentino K, Bradbury D, Cosentino M, Diaz-Mayoral N, Kennedy M, Engel T, Williams P, Erickson K, Ardlie K, Winckler W, Getz G, DeLuca D, MacArthur D, Kellis M, Thomson A, Young T, Gelfand E, Donovan M, Meng Y, Grant G, Mash D, Marcus Y, Basile M, Liu J, Zhu J, Tu Z, Cox N J, Nicolae D L, Gamazon E R, Im H K, Konkashbaev A, Pritchard J, Stevens M, Flutre T, Wen X, Dermitzakis E T, Lappalainen T, Guigo R, Monlong J, Sammeth M, Koller D, Battle A, Mostafavi S, McCarthy M, Rivas M, Maller J, Rusyn I, Nobel A, Wright F, Shabalin A, et al 2013 The Genotype-Tissue Expression (GTEx) project *Nat. Genet.* **45** 580–5

[36] Anon Surveillance, Epidemiology , and End Results Program. National Institute of Health (NIH) - Cancer Stat Facts: Female Breast Cancer Subtypes

[37] Anon Surveillance, epidemiology, and end results program (SEER)

[45] Zhu C, Kong Z, Wang B, Cheng W, Wu A and Meng X ITGB3/CD61: a hub modulator and target in the tumor microenvironment

[61] Anon Expression of MAPK6 in cancer - Summary - The Human Protein Atlas

[62] Takahashi C, Miyatake K, Kusakabe M and Nishida E 2018 The atypical mitogen-activated protein kinase ERK3 is essential for establishment of epithelial architecture *J. Biol. Chem.* **293** 8342–61

[63] Elkhadragy L, Alsaran H, Morel M and Long W 2018 Activation loop phosphorylation of ERK3 is important for its kinase activity and ability to promote lung cancer cell invasiveness *J. Biol. Chem.* **293** 16193–205

[64] Long W, Foulds C E, Qin J, Liu J, Ding C, Lonard D M, Solis L M, Wistuba I I, Qin J, Tsai S Y, Tsai M-J and O'Malley B W 2012 ERK3 signals through SRC-3 coactivator to promote human lung cancer cell invasion *J. Clin. Invest.* **122** 1869–80

[65] Bogucka K, Marini F, Rosigkeit S, Schloeder J, Jonuleit H, David K, Schlackow M and Rajalingam K 2020 ERK3/MAPK6 is required for KRAS-mediated NSCLC tumorigenesis *Cancer Gene Ther.*

[66] Quinn H M, Vogel R, Popp O, Mertins P, Lan L, Messerschmidt C, Landshammer A, Lisek K, Château-Joubert S, Marangoni E, Koren E, Fuchs Y and Birchmeier W 2021 YAP and β-Catenin Cooperate to Drive Oncogenesis in Basal Breast Cancer *Cancer Res.* **81** 2116–27

[67] Luo J, Zou H, Guo Y, Tong T, Chen Y, Xiao Y, Pan Y and Li P 2023 The oncogenic roles and clinical implications of YAP/TAZ in breast cancer *Br. J. Cancer* **128** 1611–24

[68] Xu L, Corcoran R B, Welsh J W, Pennica D and Levine A J WISP-1 is a Wnt-1- and [illegible]-catenin-responsive oncogene

[69] Huang Y-T, Lan Q, Lorusso G, Duffey N and Rüegg C 2017 The matricellular protein CYR61 promotes breast cancer lung metastasis by facilitating tumor cell extravasation and suppressing anoikis *Oncotarget* **8** 9200–15

[70] Espinoza I, Kurapaty C, Park C-H, Steen T V, Kleer C G, Wiley E, Rademaker A, Cuyàs E, Verdura S, Buxó M, Reynolds C, Menendez J A and Lupu R Depletion of CCN1/CYR61 reduces triple-negative/basal-like breast cancer aggressiveness

**Table 1. qPCR primers for gene expression determination**

| Gene name | Protein name | Forward primer 5’-3’ | Reverse Primer 5’-3’ |
|---|---|---|---|
| MAPK6 | ERK3 | GGTCTTGCACGGATCATGGA | GTGCACCTGCAAAAAGGGTT |
| YAP1 | YAP | AGAAGAGGTACCATGCTGTCCCAGATGAACGTCACA | ACAACATCTAGAATCCCGGGAGAAGACACTGGATTT |
| CYR61 | CYR61 | CCCGTTTTGGTAGATTCTGG | GCTGGAATGCAACTTCGG |
| KRT14 | Cytokeratin 14 | TGAGCCGCATTCTGAACGAG | GATGACTGCGATCCAGAGGA |
| SNAI1 | SNAIL | TCGGAAGCCTAACTACAGCGA | AGATGAGCATTGGCAGCGAG |
| CTNNB1 | Beta-catenin 1 | AAAGCGGCTGTTAGTCACTGG | CGAGTCATTGCATACTGTCCAT |
| ACTB | Beta-actin | TGACGTGGACATCCGCAAAG | CTGGAAGGTGGACAGCGAGG |
